# GAS7 coordinates the activity of RAC1 and CDC42 to enhance macropinocytic surveillance and suppress motility in macrophages

**DOI:** 10.64898/2025.12.09.693239

**Authors:** Anaël Hanouna, Mabel San-Roman, Victorine Hilly, Sarah Taheraly, François-Xavier Gobert, Mathieu Maurin, Camille Petit, Emma Granier, Philippe Benaroch, Vasco Rodrigues

## Abstract

Fluid uptake by macropinocytosis is central for the sentinel function of macrophages. However, how the actin network is organized at the cell surface to drive persistent membrane ruffling and macropinocytic cup formation remains unclear. Due to the high curvature of dorsal ruffles, we asked whether membrane curvature-associated proteins, such as those containing F-BAR domains, participate in this process. Here, leveraging public immune transcriptomic profiling and functional assays, we identify Growth Arrest Specific-7 (GAS7) as an F-BAR protein required for membrane ruffling and macropinocytosis in primary human monocyte-derived macrophages. GAS7 is enriched at dorsal ruffles and associates with actin regulators, sustaining high levels of active RAC1, to promote ruffle assembly. Moreover, GAS7 restrains the activation of CDC42, a GTPase linked to filopodia formation, cell motility and invasion. As a result, loss of GAS7 reduces ruffling and fluid uptake and instead promotes a phenotype characterized by increased motility, prominent filopodia, altered podosome dynamics, as well as an extracellular matrix-remodelling transcriptional program. Finally, we demonstrate that GAS7 enhances innate surveillance by augmenting the inflammatory response to muramyl dipeptide (NOD2 ligand) and by modulating the kinetics of TLR4 signalling to lipopolysaccharide. Thus, by coordinating RAC1 and CDC42 activities, GAS7 promotes macropinocytic uptake and enhances the macrophage responsiveness to microbial cues, while suppressing a more motile and invasive program.

## Introduction

Macrophages are critical sentinels of innate immunity that continually survey tissues for signs of infection or damage(1). To fulfil this role, they internalise large volumes of extracellular fluid through macropinocytosis(2). This non-selective uptake allows them to ingest pathogens, cellular debris and aggregates for lysosomal degradation(3). Macropinocytosis also supplies microbial ligands, allowing efficient activation of innate receptors and amplification of the immune response(2,3).

Morphologically, macropinocytosis begins when the plasma membrane protrudes outward to form ruffles. These ruffles give rise to macropinocytic cups whose tips fuse with each other giving rise to large vacuoles known as macropinosomes(4). Macropinosomes then mature through the endolysosomal system where cargoes are digested(5). While many cell types can macropinocytose in response to growth factors or other stimuli, macrophages and dendritic cells (DCs) are distinctive in that they perform it constitutively(2). Some tumours, notably pancreatic cancers, co-opt this constitutive uptake to scavenge extracellular proteins and sustain their growth in nutrient-poor environments(6).

The assembly of membrane ruffles and macropinocytic cups is driven by actin polymerization and requires coordinated changes in membrane lipid composition and small GTPase signalling(7). In macrophages, constitutive ruffling is triggered by extracellular Ca²⁺ which activates the calcium-sensing receptor (CaSR), and initiates signalling cascades leading to activation of the RAC1 GTPase(8–12). RAC1 subsequently engages effectors such as the WAVE complex to promote Arp2/3-mediated branched actin polymerization within ruffles(13). To convert a ruffle into a sealed macropinosome, RAC1-mediated signalling at the edge of the cup must be attenuated to halt actin polymerization(11,14,15). This allows sealing of the macropinocytic cup and formation of a nascent macropinosome that matures through the endocytic pathway.

Despite considerable progress in identifying components of the macropinocytic machinery, little is known about how cytoskeletal modulators are targeted to the membrane and retained there to sustain local actin polymerisation and ruffle formation. Curvature is a key morphological feature of membrane ruffles, suggesting that curvature-associated proteins help regulate this process. The BAR superfamily, which includes proteins with BAR, F-BAR and I-BAR domains, are membrane curvature-associated proteins that couple membrane dynamics to actin remodelling during processes such as migration, endocytosis and autophagy(16,17). In the amoeba *Dictyostelium discoideum*, the BAR-domain protein RGBARG orchestrates the assembly of macropinocytic cups by coordinating the location of active RAS and RAC1. However, no analogous factors have been identified in macrophages, limiting our understanding of how membrane shape regulates macropinocytosis in immune cells.

Another layer of complexity arises from the apparent antagonism between macropinocytosis and cell migration(18,19). Immature DCs engage in constitutive macropinocytosis but switch it off when they mature and migrate to lymph nodes(20). In macrophages, this dichotomy is less clear, although cells moving along chemotactic gradients take up less fluid than resting sentinels(21). Both migration and macropinocytosis require the activity of Rho-family GTPases. The RAC1/WAVE axis is indispensable for macropinocytosis, but also drives lamellipodial protrusions that propel cell migration(22). It remains unclear how the activation of RAC1 is directed towards the formation membrane ruffles or lamellipodia. CDC42, another important Rho family GTPase, has disputed roles in macropinocytosis in myeloid cells (7,23), but it is essential for filopodia formation(24), and for podosome assembly that enable macrophages to traverse the extracellular matrix (ECM)(25,26). How GTPase activation in macrophages is regulated to ensure efficient ruffling and macropinocytic uptake, while repressing movement and ECM-remodelling, remains unclear.

Here, we identify Growth Arrest Specific-7 (GAS7), a curvature-sensing protein of the F-BAR family, as a key regulator of membrane ruffling in primary human macrophages. GAS7 has been previously implicated in phagocytosis in mouse macrophages (27) and as a regulator of metastatic invasion by tumour cells (28–30). We show that, in human MDMs, GAS7 localizes to membrane ruffles where it binds several cytoskeletal regulators and sustains elevated RAC1 activity, thereby promoting fluid uptake and enhancing macrophage responses to microbial ligands. In its absence, ruffling is severely reduced and macrophages acquire a distinct phenotype marked by prominent filopodia, altered podosome dynamics, increased motility and expression of ECM-remodelling factors. Our data indicates that these alterations arise from chronic aberrant activation of CDC42 when GAS7 is absent. By modulating the balance between RAC1 and CDC42 signalling, GAS7 enhances the sentinel functions of macrophages and clarifies how cells prioritize between fluid uptake versus movement.

## Results

### GAS7 is an F-BAR protein required for optimal macropinocytosis in macrophages

Macrophages continuously form curved membrane ruffles, suggesting that curvature-associated proteins play a role in their assembly or dynamics. To identify BAR domain–containing proteins enriched in myeloid cells, we mined public single-cell RNA-seq datasets (31)(32)(33) and found that the F-BAR family members *GAS7*, *FES*, and *SRGAP1* are significantly enriched in myeloid populations among immune cell types (Fig. S1a-c). GAS7 also ranked among the most highly expressed BAR domain–containing genes in our bulk RNA-seq data from human monocyte-derived macrophages (MDMs; Fig. S1d), prompting us to examine its role in macropinocytosis.

In humans and mice, GAS7 exists as three isoforms generated by alternative splicing. These isoforms differ in their N-terminal scaffolding domains (SH3 and WW) but all share a C-terminal F-BAR domain (Fig. S1e). While fresh human monocytes express only isoform a, differentiated MDMs express all three, with isoform c predominating (Fig. 1a, S1f). Confirming the public transcriptomic data, we found that GAS7 is constitutively expressed in additional myeloid cells such as conventional DC1s (cDC1) and cDC2s but is absent from non-myeloid immune cells including CD4⁺ T cells and plasmacytoid DCs (Fig. 1a, S1f).

**Figure 1.**
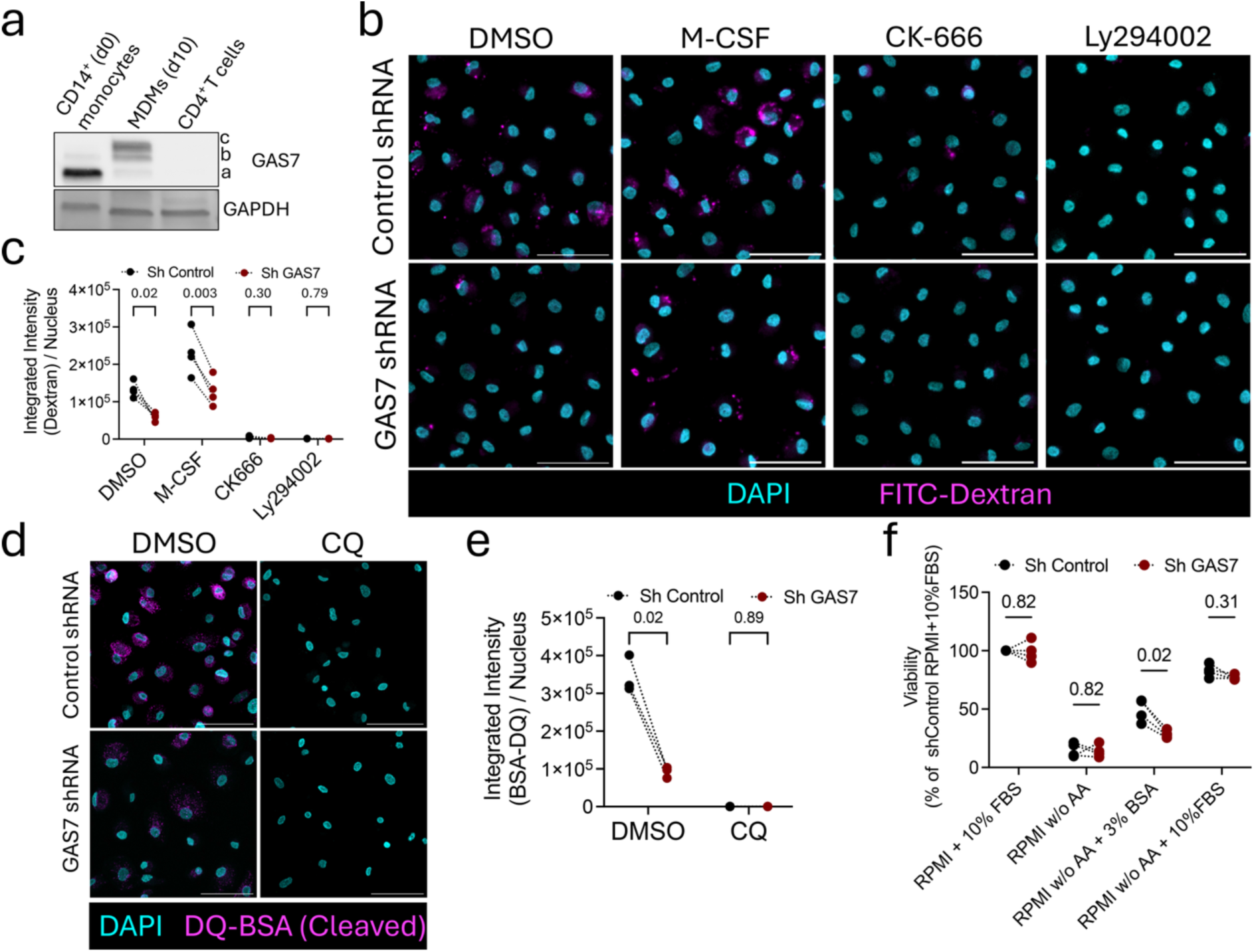
GAS7 is required for optimal macropinocytosis in human macrophages. **a.** Western Blot depicting the expression of GAS7 in fresh human monocytes, 7-day differentiated MDMs and CD4^+^ T cells from the same donor. **b.** Representative confocal images (SUM z-projections) of fixed shControl or shGAS7 MDMs exposed to 70 Kda FITC-Dextran (200µg/mL) for 30 min and subjected to the indicated treatments. M-CSF (200 ng/mL) was added simultaneously with dextran, while for DMSO (0.1% V/V), CK-666 (20µM) and LY294002 (10µM), cells were pretreated for 1 hour with inhibitors before addition of dextran. Scale bars = 50µm. **c.** Quantification of dextran uptake for 4 independent donors. The total integrated intensity for the dextran channel was quantified from an acquired field and divided by the number of nuclei in the field. Each symbol represents the average dextran signal per nuclei for each donor (N=4). 300 to 500 cells were acquired per condition per donor. **d.** Representative confocal images (SUM z-projection) of fixed shControl or shGAS7 MDMs exposed to DQ-Red BSA (10µg/mL) for 3 hours in non-treated (DMSO 0.1% V/V) or chloroquine (CQ, 50µM)-treated conditions. Scale bars = 50µm. **e.** Quantification of DQ-BSA uptake and cleavage from (d) for 3 independent donors. The total integrated intensity for the DQ-BSA channel was quantified from an acquired field and divided by the number of nuclei in the field. Each symbol represents the average DQ-BSA signal per nuclei for each donor (N=3). 100-200 cells were acquired per condition per donor. **f.** Viability, determined by the CellTitter-Glo® assay, of shControl or shGAS7 MDMs cultured in the indicated conditions for 10 days. Each symbol represents an independent donor (N=4). Values were normalized relative to the 4-donor average of shControl cells cultured in RPMI + 10% FBS (100%). Paired t test between shControl and shGAS7 for each culture condition. P-value of paired t-test between shControl and shGAS7 for each condition is indicated above brackets.

To investigate the role of GAS7 in macropinocytosis, we silenced its expression in primary human MDMs using lentiviral transduction of shRNA (Fig. S1g) and measured uptake of fluorescent 70-kDa dextran after a 30-min pulse. GAS7-silenced cells internalized significantly less dextran at baseline and following M-CSF stimulation (Fig. 1b–c). As expected, dextran uptake was inhibited by the Arp2/3 inhibitor CK-666 and the PI3K inhibitor LY294002 (Fig. 1b–c). Conversely, lentiviral overexpression of GAS7c (Fig. S1h) increased dextran uptake relative to a control vector (Fig. S1i-j). Because macropinocytosed proteins traffic to the endolysosomal system for degradation, we used DQ-Red BSA to assess degradative uptake. GAS7-silenced macrophages showed reduced levels of cleaved, dequenched DQ-BSA after a 3-h exposure, confirming their decreased capture of extracellular protein (Fig. 1d–e). In RAS-transformed tumour cells, macropinocytosis supports cell survival under amino-acid deprivation by promoting the scavenging of extracellular proteins (34). Similarly, we observed that the expression of GAS7 conferred a survival advantage to MDMs cultured for 10 days without free amino acids, and relying on extracellular BSA as amino acid source, whereas under standard cell culture conditions, the absence of GAS7 had no impact on MDM viability (Fig. 1f). Although clathrin-mediated endocytosis (CME) also depends on actin polymerization(35), GAS7 did not affect transferrin uptake, a canonical CME marker (Fig. S1k), indicating that it is not associated with all endocytic pathways. Collectively, these data identify GAS7 as a macrophage-enriched F-BAR protein required for optimal macropinocytic flux and led us to assess where GAS7 acts within the macropinocytic pathway.

### GAS7 localizes to membrane ruffles in macrophages

We examined the subcellular localization of GAS7 in human MDMs. Several commercial antibodies failed to detect endogenous GAS7 in MDMs by immunofluorescence. To circumvent this, we transduced MDMs with EGFP-tagged GAS7c driven by a *PGK* promoter to achieve near-endogenous expression levels (Fig. S2a, Video S1). We also expressed a shRNA-resistant EGFP-GAS7C construct in GAS7-silenced MDMs, also driven by the *PGK* promoter (Video S2). In fixed and live confocal imaging, EGFP–GAS7c was enriched at membrane ruffles, although a diffuse cytoplasmic signal was also visible (Fig. 2a, 2c; Videos S1-2). Fractionation of MDM lysates confirmed that endogenous GAS7c is distributed between membrane and cytosolic fractions, validating the EGFP constructs (Fig. S2b). EGFP–GAS7c colocalized with F-actin at the dorsal (ruffle-enriched) surface but not with actin-rich podosomes at the ventral surface (Fig. 2a-b). It also colocalized at the surface with WAVE2, a ruffle marker in macrophages (36) (Fig. 2c-d).

**Figure 2.**
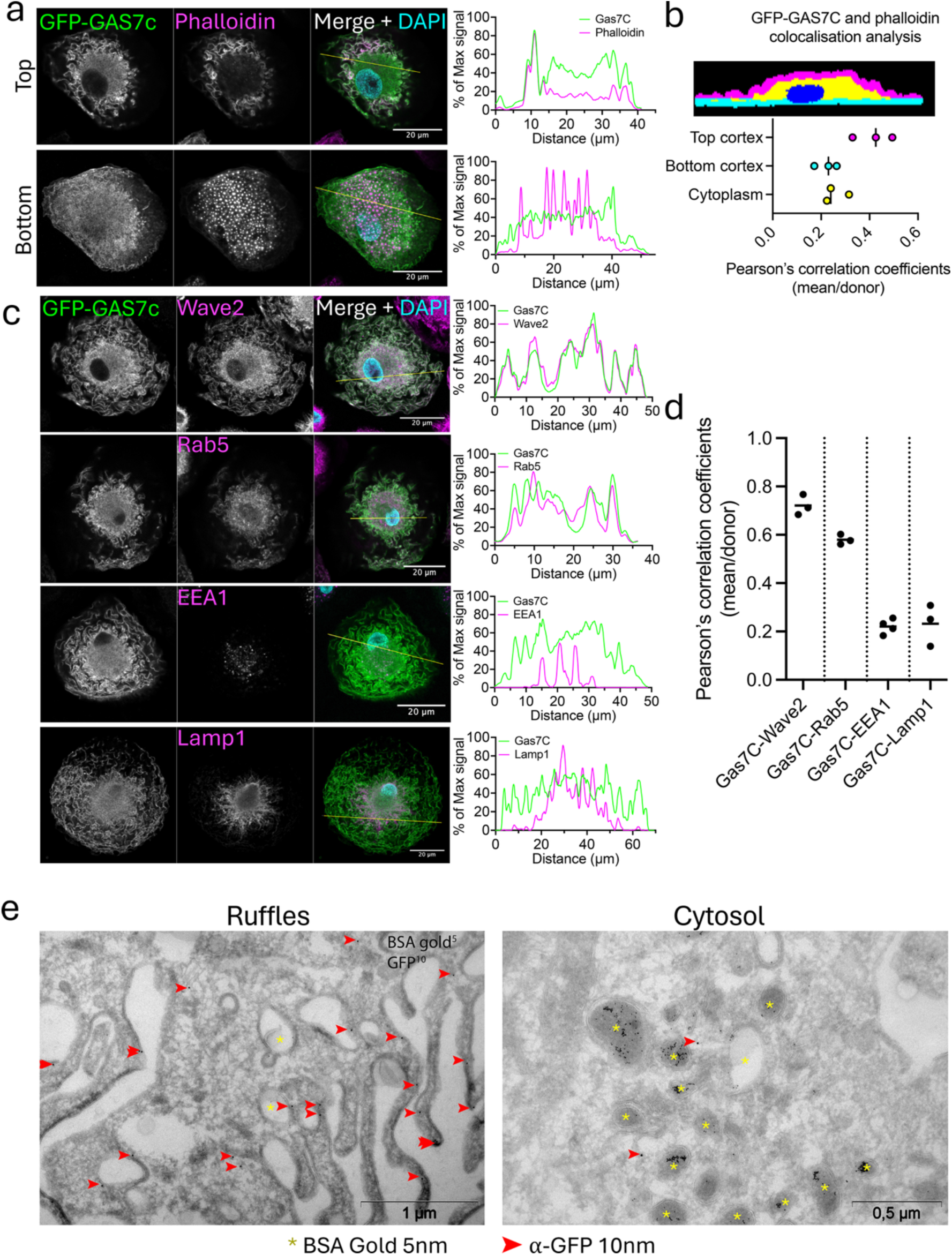
GAS7 localises in membrane ruffles in macrophages. **a.** Representative confocal images of MDMs transduced with EGFP-GAS7C and stained with phalloidin. Z-sections (thickness =0.31µm) at the top, dorsal part of the cell and the bottom, ventral part are depicted. The graph on the right shows the intensities of both EGFP-GAS7C and phalloidin signals plotted along the yellow line for both dorsal (top, ruffles) and ventral (bottom, podosomes) sections. Intensity is represented as a percentage of the maximum signal of the staining and smoothed with the 8 closest neighbours. **b.** Pearson’s correlation coefficient (PCC) between the pixel intensities for EGFP-GAS7C and Phalloidin staining. As exemplified in the cell mask above, MDMs were segmented into cytoplasm (yellow), ventral cortex (cyan) and dorsal cortex (magenta) and PCCs were assessed independently for each cell section. Each dot represents the mean PCC for an independent donor (N=3), calculated from at least 5 GFP^+^ cells per donor. **c.** Representative confocal images of MDMs transduced with EGFP-GAS7C and immunostained for either Wave2, Rab5, EEA-1 or Lamp1. The images are from a single Z-section (thickness =0.31µm) at the dorsal part of the cell. On the right the intensities of both GFP-GAS7c and the indicated marker are plotted along the yellow line. Intensity is represented as a percentage of the maximum signal of the staining and smoothed with the 10 closest neighbours. **d.** Mean PCC calculated between EGFP-GAS7C and the co-staining, for each donor (N=3-4, as indicated). Values are calculated on the total volume of the cell (excluding the nucleus). Representative images of at least 5 GFP+ cells per donor. **e.** MDMs were transduced with EGFP-GAS7c, sorted for GFP+ cells, and exposed to BSA coupled to gold particles of 5-nm for 2 hours. Ultrathin cryosections were labelled with an antibody specific for EGFP and Protein A coupled to gold particles of 10 nm diameter. Representative images from one donor of 2 donors processed.

In contrast, GAS7c showed only partial colocalization with the early macropinosome marker Rab5 and was absent from EEA1⁺ early endosomes and LAMP1⁺ late endosomes/lysosomes (Fig. 2c-d), suggesting it disengages from the membrane after macropinosome closure. In agreement, live-cell imaging of EGFP–GAS7c-expressing MDMs exposed to CF633-dextran confirmed its dynamic enrichment at ruffles and absence from dextran-filled macropinosomes as they appear in the cytoplasm (Video S3). Finally, immuno-electron microscopy of macrophages expressing EGFP–GAS7c, and pulsed with BSA–gold particles, confirmed the localization of GAS7 in membrane ruffles with ultrastructural resolution, but not to BSA-containing macropinosomes (Fig. 2e). Thus, GAS7 concentrates at dorsal ruffles but disengages from the membrane before macropinosome maturation, suggesting a role in ruffle assembly/dynamics rather than endosomal trafficking. We therefore tested whether GAS7 is required for ruffle formation.

### GAS7 sustains membrane ruffling and restrains the transition to a motile phenotype in macrophages

To explore a possible role for GAS7 in the formation of membrane ruffles we examined scanning electron microscopy (SEM) images of MDMs. In control shRNA-transduced cells, we observed the characteristic florid ruffles that thoroughly coat the dorsal surface of the cell (Fig. 3a). In contrast, GAS7-silenced MDMs exhibited a marked reduction in ruffle density, size, and complexity, along with an apparent increase in cell spreading on the substrate (Fig. 3a). To quantify this phenotype, we cultured MDMs, transduced with shControl or shGAS7 and previously stained with cytoplasmic dies of different colours, together in the same well to minimize acquisition bias. Then, we employed fluorescent phalloidin to stain F-actin and scored the presence and magnitude of membrane ruffles in a blinded manner (Fig. 3b, Fig. S3a-c). In control cells, ∼70% displayed dense or intermediate ruffles; this fell below 30% in GAS7-silenced cells, which frequently exhibited linear, actin-rich filopodia-like projections (Fig. 3b). GAS7-silenced MDMs also showed a significantly larger ventral contact area (Fig. 3d), confirming the increased spreading apparent in SEM images.

**Figure 3.**
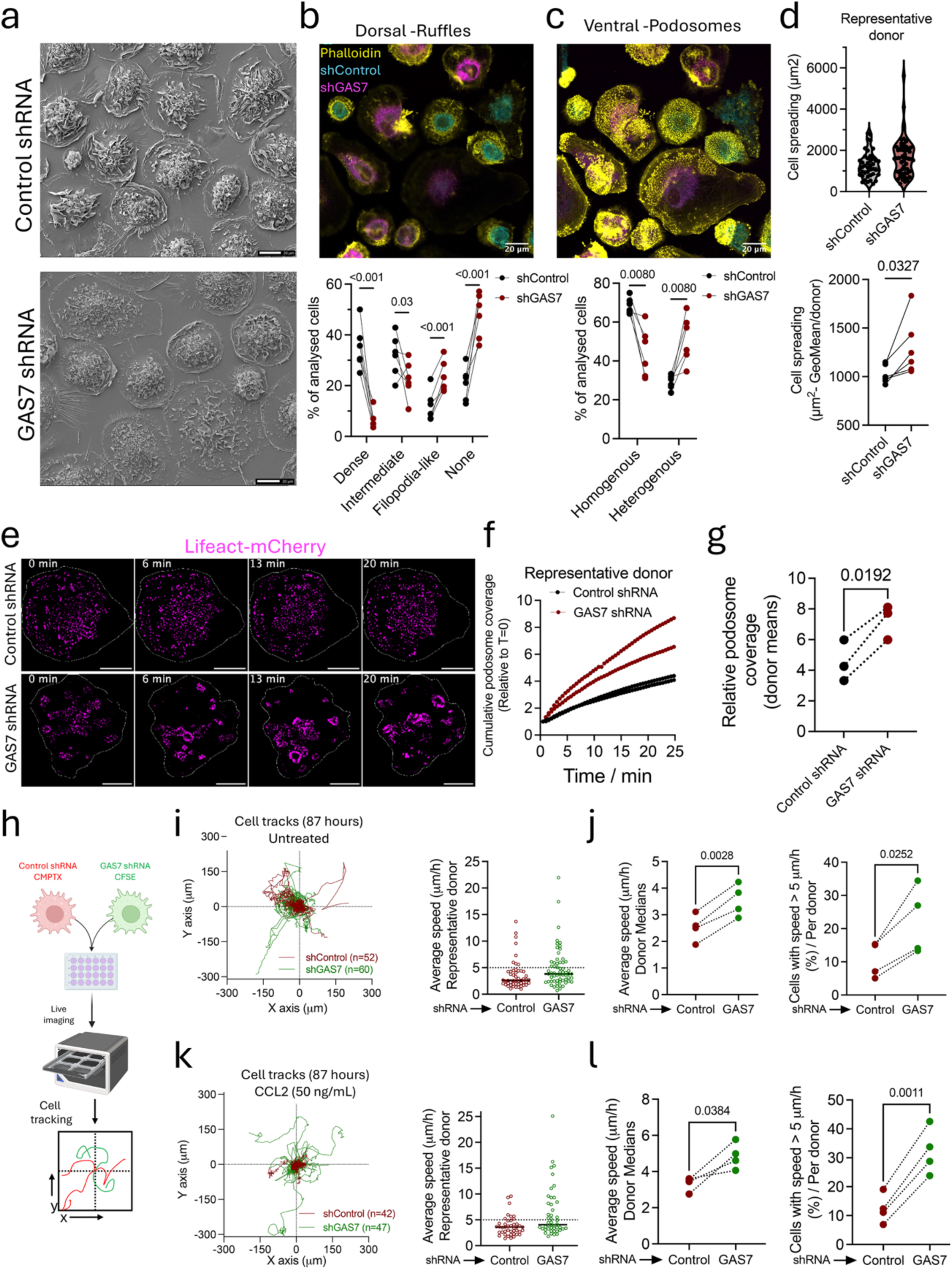
GAS7 sustains membrane ruffling and restrains the transition to a motile phenotype in macrophages. **a.** Scanning electron micrographs of shControl or shGAS7 transduced MDMs. Representative images from one of N=2 donors processed. **b-c.** Top panels depict representative images of F-actin staining of shControl (Cyan) and shGAS7 (Magenta) MDMs acquired at the dorsal (b) and ventral (c) planes of the cells (z thickness =0.31 µm). Experiments in which the cytoplasmic dyes were reversed, were also performed, and pooled for the quantifications. The plots show the quantification of membrane ruffles (b) and podosome organization (c) for N=6 donors (see Fig. S3A-B and Methods for further details). Each dot represents the percentage of cells in each category for a given donor, calculated from 30-90 acquired cells per condition per donor. Multiple paired t-test between shControl and shGAS7 cells. **d.** Ventral surface of shControl and shGAS7 MDMs, calculated from the cells analysed in b-c. The top plot shows a representative donor, each dot representing an individual cell (n=40-60 cells). The bottom panel shows the quantification for multiple donors, with each dot representing the average value for each independent donor (N=6). Paired t-test between shControl and shGAS7. **e.** Representative timeframes from live imaging of MDMs transduced with Control shRNA (top) and GAS7 shRNA (bottom) and co-transduced with Lifeact mCherry to visualize F-actin. Shown are max intensity projections of the Lifeact signal. Scale bar=20 µm. See also Video S4. **f-g.** Quantification of podosome dynamics in Lifeact-expressing shControl and shGAS7 MDMS. The left plot depicts the podosome relative cumulative coverage of the MDM ventral surface during the acquisition time for n=2 cells from a representative donor. The right plot displays the relative cumulative coverage for N=4 independent donors. See also Video S5. **h.** Schematic representation of the experimental approach used to quantify MDM movement. Cells (shControl or shGAS7) were stained with the indicated cell masks and seeded together in culture plates, in the presence or absence of CCL2 and imaged for 87 hours, at a frame acquisition rate of 1 per hour. See also Fig S3e-i for the equivalent experiment with the dyes reversed. **i.** Trajectories (left) and average speeds (right) of individual MDMs from a representative donor in the untreated condition and pre-stained as indicated in (h), over 87 hours of live imaging. The plots were built from 2 acquisition fields, including the one displayed in Video S6. Single cells are represented by individual trajectories in (left) and symbols in (right). The black horizontal line represents the population median. **j.** Median average speeds (left) and percentage of cells faster than 5 µm/h (right) for 4 independent donors, for the untreated condition. Paired t test. **k.** Trajectories (left) and average speeds (right) of individual MDMs from a representative donor in CCL2-exposed cells and pre-stained as indicated in (h), over 87 hours of live imaging. The plots were built from 2 acquisition fields, including the one displayed in Video S6. Single cells are represented by individual trajectories in (left) and symbols in (right). The black horizontal line represents the population median. **l.** Median average speeds (left) and percentage of cells faster than 5 µm/h (L) for 4 independent donors, for CCL2-treated condition. Paired t test.

Although GAS7 does not colocalize with podosomes in MDMs (Fig 2a-b), their organization was altered in its absence. Whereas most control MDMs displayed homogeneously distributed podosomes across the ventral surface, about 50% of GAS7-silenced cells exhibited polarized or heterogeneous podosome patterns (Fig. 3c, S3b). To further explore how the absence of GAS7 impacts podosome dynamics, we performed live imaging on Lifeact-mCherry-expressing MDMs, simultaneously transduced with an shControl or shGAS7 (Fig S3d). This approach confirmed that silencing GAS7 modifies podosome dynamics, as manifested by increased turnover and altered patterning. When GAS7 is silenced, podosomes often appear organized as rosette structures that exhibit fast cycles of assembly and disassembly and that cumulatively cover the ventral surface of the cell faster than in control cells (Fig. 3e-g; Videos S4-5).

Because these changes suggested a shift toward a more motile phenotype, we quantified the 2D motility of shControl and shGAS7 macrophages, cultured *in vitro* in the same well. While human MDMs are almost immotile in plastic culture plates, GAS7 silencing boosted their stochastic motility at the steady-state and in the presence of the chemokine CCL2 (Fig. 3h–m, Fig. S3e-i, Videos S6-7). Overall, GAS7 is required for optimal assembly of dorsal ruffles, and its loss promotes cell spreading and the formation of filopodia with altered podosome dynamics and increased motility.

### Macrophages express ECM-remodelling factors in the absence of GAS7

GAS7 limits macrophage motility and regulates podosome dynamics. As podosomes coordinate the secretion of matrix-remodelling enzymes and are necessary for invasion(37,38), these observations point toward a pro-invasive shift upon GAS7 depletion. Because transitions toward motile/invasive states in macrophages are typically coupled to widespread transcriptional reprogramming(39,40), we compared the transcriptome of shControl and shGAS7 MDMs from six independent donors by bulk RNA-seq (Fig. 4a–g). Transcripts involved in ECM remodelling, including *THBS1*, collagens (*COL4A1*, *COL6A1*) and laminins (*LAMB1*, *LAMB3*), were among the most upregulated in GAS7-silenced MDMs (Fig. 4a). Pathway enrichment analysis (Fig. 4b–c) and gene set enrichment analysis (GSEA) (Fig. 4d-e) confirmed a transcriptomic rewiring toward an ECM-remodelling profile. Moreover, chemokines including CXCL8 and CCL2 were upregulated at both transcript and protein levels (Fig. 4a-b, f-g), indicating a broader shift toward a pro-remodelling state. Yet, we found no evidence of general induction of a pro-inflammatory state, as the levels of *TNF*, *IL23* or *IL1A* remained stable after silencing GAS7 (Fig. S4a). Ruling out an off-target effect, a second shRNA against GAS7 (Fig S1g) also induced a spontaneous upregulation of *THBS1* and *CXCL8* (Fig. S4b). Together, these data show that GAS7 loss re-establishes transcriptional programs linked with ECM remodelling, prompting us to explore the upstream mechanisms through which GAS7 sustains ruffling and restrains invasiveness.

**Figure 4.**
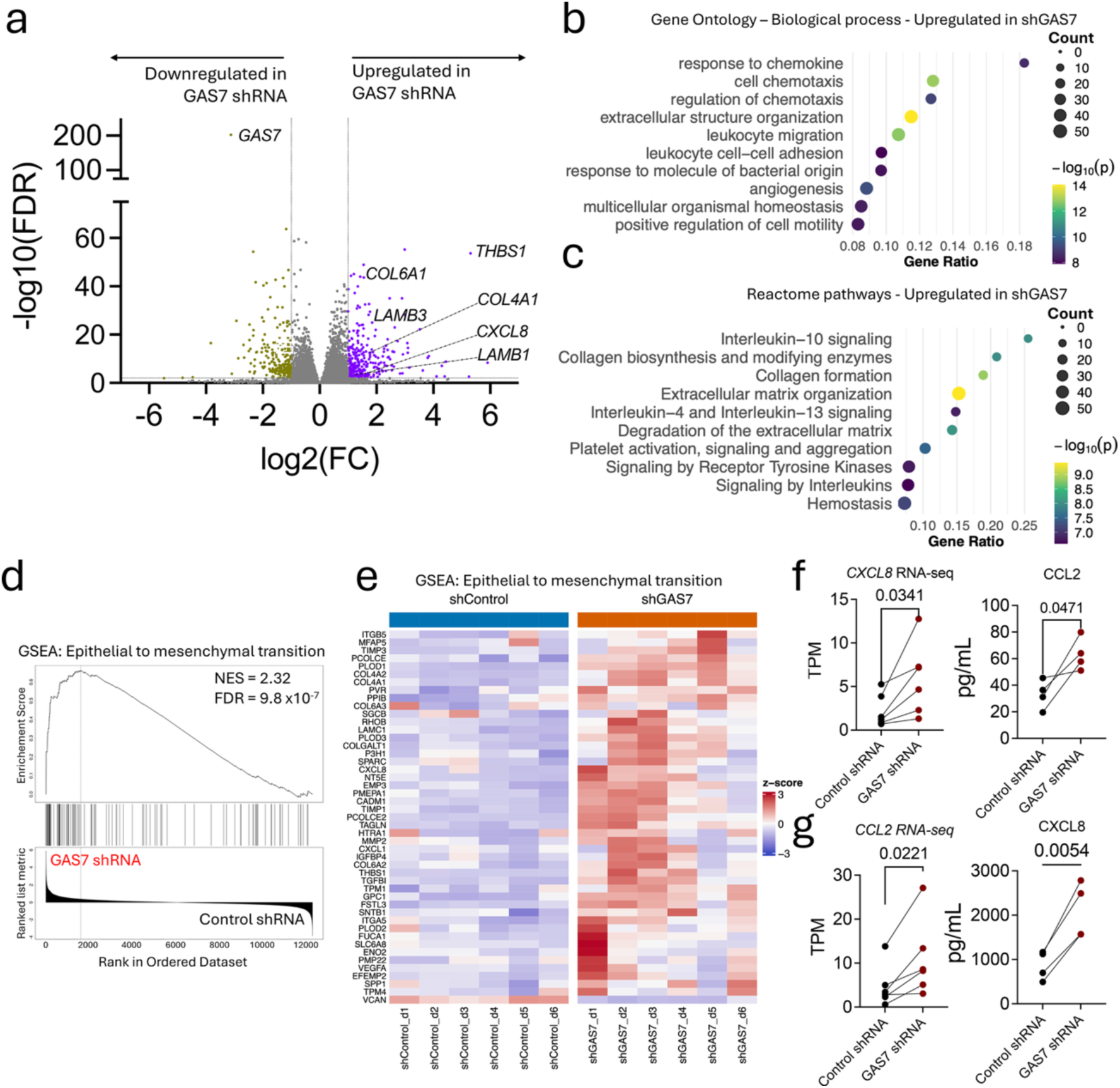
Silencing GAS7 rewires macrophages towards an ECM-remodelling transcriptome. **a.** Volcano plot depicting differentially expressed genes (DEGs) between shControl and shGAS7 MDMs. Genes were considered as differentially expressed when –log10(FDR)>2 and log2(FC)< -1 or log2(FC)>1. **b.** Top 10 Gene ontology (Biological Process) pathways ranked by gene ratio and significantly enriched in shGAS7 MDMs. **c.** Top 10 reactome pathways ranked by gene ratio and significantly enriched in shGAS7 MDMs. **d.** Enrichment plot for the Epithelial to Mesenchymal Transition (EMT) pathway; the top enriched GSEA pathway in shGAS7 MDMs. The black curve is the running enrichment score, showing the accumulation of EMT genes as ranked by their differential expression. Black vertical lines denote the positions of EMT pathway genes in the ranked gene list. **e.** Heat-map of the top ranked EMT genes between shControl and shGAS7 MDMs. **f.** Expression of CXCL8 in shControl and shGAS7 MDMs in the RNAseq dataset (left) and spontaneous secretion of CXCL8 to the supernatant in shControl and shGAS7 MDM cultures. Each dot is an independent donor. Paired t test. **g.** Expression of CCL2 in shControl and shGAS7 MDMs in the RNAseq dataset (left) and spontaneous secretion of CCL2 to the supernatant in shControl and shGAS7 MDM cultures. Each dot is an independent donor. Paired t test. FDR = False discovery rate; FC = Fold change; NES =Normalized Enrichment Score; TPM = transcripts per million

### GAS7 binds subunits of the WAVE complex and sustains RAC1 activation

To explore the mechanistic basis for the phenotypes observed in the absence or presence of GAS7 in MDMs, we identified its binding partners through a yeast two-hybrid (Y2H) screen using full-length GAS7c as bait against a human MDM cDNA library (Fig. 5a, S5a). Among the high-confidence hits were several cytoskeleton-associated proteins (Fig. 5a, S5a–b), including previously reported GAS7 partners, such as GAS7 itself(28,41,42). Notably, the screen recovered ABI1 and WAVE2, both subunits of the WAVE regulatory complex (WRC) which activates Arp2/3 to promote actin polymerization and ruffle formation(43). Consistent with this, WAVE2 colocalized with GAS7c in ruffles (Fig. 2c–d). Activation of the WRC requires the binding of active RAC1 to relieve the complex from autoinhibition(13). In macrophages, RAC1 can be activated when extracellular Ca²⁺ engages CaSR(9,44), but additional unknown mechanisms likely sustain constitutive RAC1 activity to support continuous ruffling.

**Figure 5.**
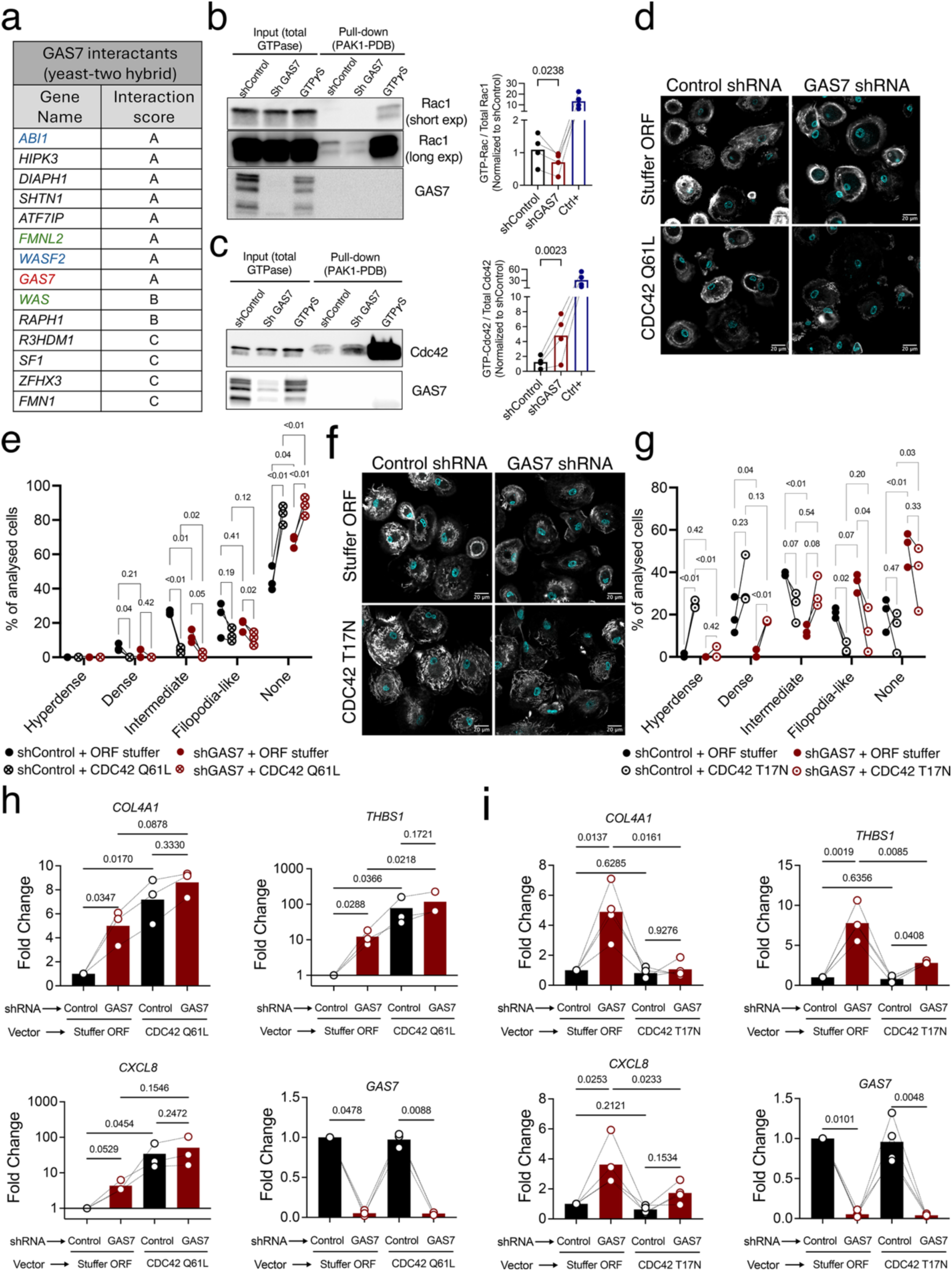
GAS7 coordinates the activation of RAC1 and CDC42 to sustain membrane ruffles and repress an ECM-remodelling phenotype. **a.** Hits retrieved from the yeast-two-hybrid screening for binding partners of GAS7 in MDMs. In blue are hits that belong to the WAVE complex, while in green are the Cdc42 effectors. GAS7 in red. The PBS (Predicted Biological Score) categorizes confidence in the interaction from very high confidence (A) to low confidence (D). See also figure S5a for more detailed information on the screen hits. **b.** GTP-RAC1 was pulled down from shControl or shGAS7 MDMs using a GST-fused PAK1-PDB and a glutathione agarose resin and analyzed by western blot. The positive control consists of shControl MDM lysates treated with an excess of GTP*γ*S. The plot on the right shows band density quantifications for theGTP-RAC1/ total-RAC1 ratio for 4 independent donors. **c.** GTP-Cdc42 was pulled down from shControl or shGAS7 MDMs using a GST-fused PAK1-PDB and a glutathione agarose resin and analysed by western blot. The positive control consists of shControl MDM lysates treated with an excess of GTP*γ*S. The plot on the right shows band density quantifications for the GTP-CDC42/ total-CDC42 ratio for 4 independent donors. **d.** Representative images of F-actin staining of shControl or shGAS7 MDMs and co-transduced with either a Stuffer ORF or CDC42(Q61L) acquired at the dorsal plane of the cells (z thickness =0.31 µm). **e.** Quantification (see Methods and Figure S3B for details) of membrane ruffles for N=6 donors for the cell types indicated. Each dot represents the average value for an independent donor, calculated from 30-90 acquired cells per condition. Multiple paired t-test between sh control and shGAS7 cells. **f.** Representative images of F-actin staining of shControl or shGAS7 MDMs and co-transduced with either a Stuffer ORF or CDC42(T17N) acquired at the dorsal plane of the cells (z thickness =0.31 µm). **g.** Quantification of membrane ruffles for N=6 donors for the cell types indicated. Each dot represents the average value for an independent donor, calculated from 30-90 acquired cells per condition. Multiple paired t-test between shControl and shGAS7 cells. **h.** qPCR for *COL4A1*, *THBS1*, *CXCL8* and *GAS7* in shControl or shGAS7 MDMs co-transduced with either a stuffer ORF or CDC42(Q61L). Each dot represents an independent donor, and the black line connects each independent donor across the different conditions. One-way ANOVA followed by Tukey’s correction for multiple comparisons. **i.** qPCR for *COL4A1*, *THBS1*, *CXCL8* and *GAS7* in shControl or shGAS7 MDMs co-transduced with either a stuffer ORF or CDC42(T17N). Each dot represents an independent donor, and the black line connects each independent donor across the different conditions. One-way ANOVA followed by Tukey’s correction for multiple comparisons.

We evaluated a role for GAS7 in sustaining RAC1 activity by pulling down GTP-bound RAC1 in GAS7-proficient and GAS7-deficient MDMs, using the PAK1 p21-binding domain. Total RAC1 levels were unchanged, but GAS7-expressing cells consistently showed higher RAC1-GTP across 5 independent donors (Fig. 5b), indicating that GAS7 sustains RAC1 activity. These data indicate that GAS7 acts as an adaptor that assembles and activates actin machinery in membrane ruffles, providing a mechanism for constitutive ruffling. As GAS7 loss led to the formation of filopodia and altered the patterns of podosome distribution and cell movement, we next asked whether CDC42 is reciprocally impacted by GAS7.

### GAS7 inhibits CDC42 activation and represses an ECM-remodelling phenotype in macrophages

Beyond WRC components, the Y2H screen indicated that GAS7 binds WASP, a CDC42 effector involved in podosome formation and dynamics(23,38,45). It also indicates that GAS7 binds FMNL2, another CDC42 effector regulating filopodia assembly, podosome function, and ECM degradation in macrophages(46) (Fig. 5a, S5a–b). We therefore measured CDC42 activity and found elevated GTP-bound CDC42 in GAS7-silenced MDMs despite unchanged total levels of CDC42 (Fig. 5c). Thus, GAS7 sustains the activation of RAC1, while repressing that of CDC42.

The role of CDC42 in membrane ruffling is unclear(7), with studies yielding conflicting results(23,47), possibly due to the partial overlap between the downstream effectors of CDC42 and those of RAC1(48). To clarify this, we tested the functional consequences of enforced CDC42 activation in human MDMs. Expression of constitutively active CDC42(Q61L) abolished membrane ruffles irrespective of GAS7 status (Fig. 5d–e) and induced extreme spreading over the substrate (Fig. S5C, Video S8). Conversely, a dominant-negative CDC42(T17N) mutant increased baseline ruffling in both control macrophages and GAS7-silenced cells (Fig. 5F–G). This confirms that CDC42 activation in human macrophages opposes ruffle formation; however, inhibition of Cdc42 alone could not completely restore membrane ruffles observed in the presence of GAS7.

While CDC42 directly targets cytoskeleton effectors, it also induces changes in gene expression by triggering signalling pathways that activate transcription factors such as NF-κB, or AP-1(49–51). In several cell types, the activation of CDC42 induces the expression of matrix metalloproteases, integrins and other factors involved in ECM remodelling (52–54). We thus hypothesized that the ECM-remodelling transcriptome observed in shGAS7 MDMs (Fig 4a-e) results from aberrant activation of CDC42. We observed that transduction of CDC42(Q61L) augmented the expression of *CXCL8*, *THBS1* and *COL4A1* (Fig. 5h), genes similarly upregulated in GAS7-silenced MDMs, while genes whose expression is unaffected by GAS7 knockdown (*TNF*, *IFI44L*) similarly remained unchanged in CDC42(Q61L) (Fig. S5d). Conversely, expressing the dominant negative CDC42(T17N) mutant significantly decreased the upregulation of *CXCL8*, *THBS1* and *COL4A1* observed upon GAS7 silencing (Fig. 5i, S5e), indicating that the transcriptional alterations observed in the absence of GAS7 are driven by aberrant CDC42 activation.

We conclude that GAS7 regulates a functional antagonism between RAC1 and CDC42 in macrophages. By promoting RAC1 activity while restraining CDC42, GAS7 sustains ruffling and prevents the adoption of a transcriptional and morphological phenotype characterized by increased motility and production of ECM-remodelling factors.

### The expression of GAS7 modulates the macrophage response to microbial ligands

Macropinocytosis contributes to immune surveillance by delivering extracellular microbial ligands to intracellular receptors(3). For example, macropinocytosis is required for the uptake and inflammatory responses to muramyl dipeptide (MDP), which activates the cytosolic NOD2 receptor(9). We thus evaluated a role for GAS7 in modulating the inflammatory response of MDMs to MDP. GAS7-silenced MDMs displayed reduced induction of *CXCL8* and *IL6* mRNAs (Fig. S6a) and diminished secretion of CXCL8, TNF and IL-6 proteins upon MDP exposure (Fig. 6a). Pharmacologic inhibition with LY294002 blunted the responses, irrespective of the expression of GAS7, supporting a macropinocytic route for the delivery of MDP to NOD2 (Fig. 6a, S6a). Thus, GAS7 is required for efficient macropinocytic delivery of NOD2 ligands.

**Figure 6.**
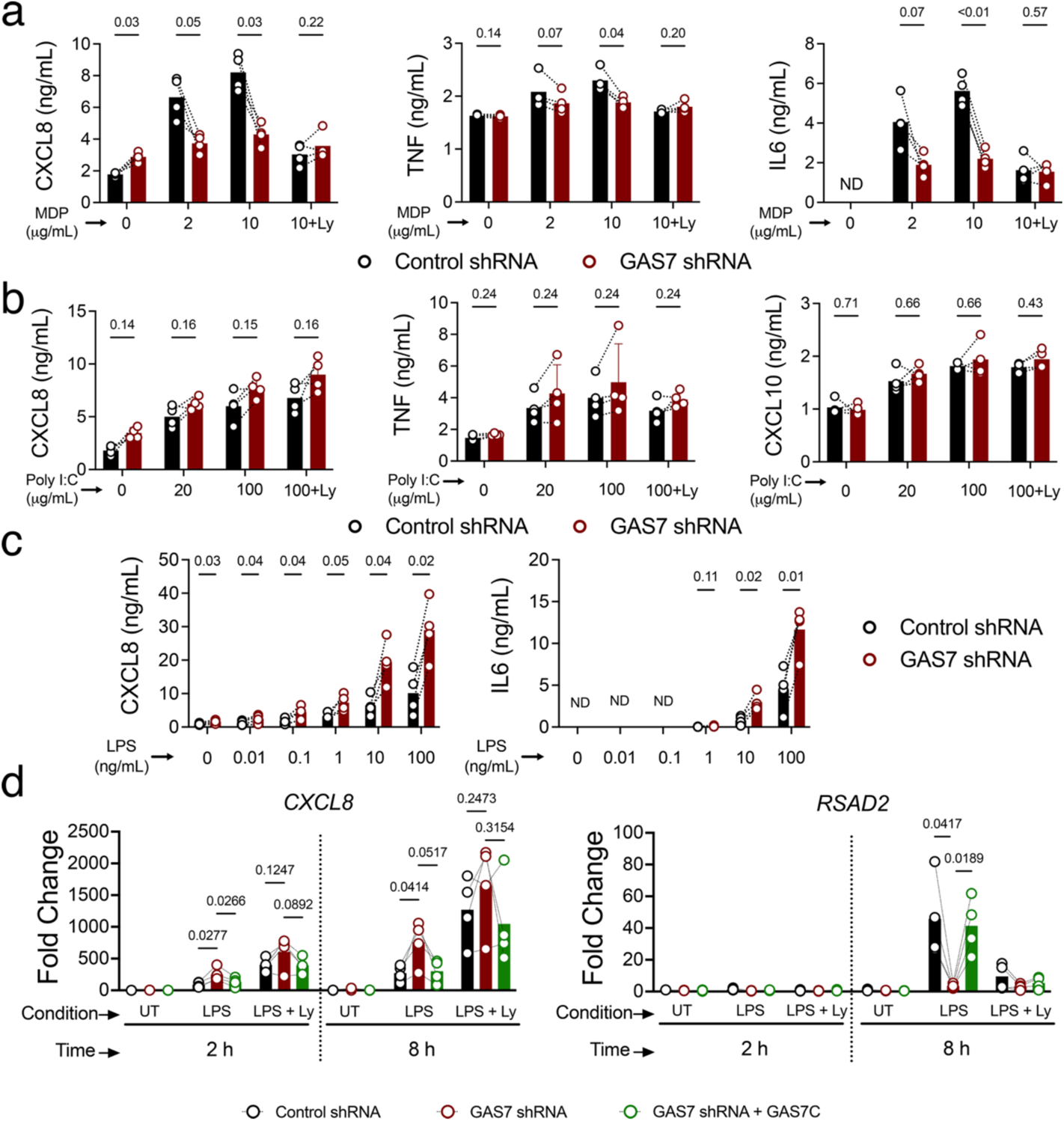
The expression of GAS7 alters macrophage response to microbial ligands. **a.** Bead-based immunoassay quantification of CXCL8, TNF and IL6 in culture supernatants from shControl or shGAS7 MDMs exposed to MDP for 16 hours, at the indicated doses, and pre-treated or not with Ly294002 (10µM). **b.** Bead-based immunoassay quantification of CXCL8, TNF and CXCL10 in culture supernatants from shControl or shGAS7 MDMs exposed to high molecular weight Poly I:C for 16 hours, at the indicated doses, and pre-treated or not with Ly294002 (10µM). **c.** Bead-based immunoassay quantification of CXCL8 and IL6 in culture supernatants from shControl or shGAS7 MDMs exposed to LPS for 16 hours, at the indicated doses. **d.** qPCR for *CXCL8* and *RSAD2* from shControl or shGAS7 MDMs exposed to LPS for 2 or 8 hours, at 100 ng/mL, and pre-treated or not with Ly294002 (10µM). Each dot represents an independent donor, and the black line connects each independent donor across the different conditions. Two-way ANOVA followed by Tukey’s correction for multiple comparisons. Each dot represents an individual donor. Multiple paired t test between shControl and shGAS7, unless indicated otherwise.

In contrast with MDP, the delivery of the double-stranded RNA mimetic poly I:C to its receptor TLR3 in endosomes occurs via a clathrin-dependent mechanism(55), while macropinocytosis appears dispensable (56). In agreement, we found that the expression of GAS7 had no impact on the induction of pro-inflammatory mediators and ISGs upon exposure to poly I:C (Fig 6b, S6b). Moreover, PI3K inhibition also did not impact the MDM response to poly I:C, confirming that it does not require macropinocytosis (Fig 6b, S6b).

Activation of TLR4 by bacterial lipopolysaccharide (LPS) initiates an early response via TIRAP/Myd88 from the plasma membrane leading to rapid NF-κB–dependent induction of inflammatory mediators such as IL6 and CXCL8(57). A second wave of the response to LPS requires TLR4 internalization and signals through the adaptors TRAM/TRIF to induce type I IFN and ISGs(58). Evidence suggests that, among other internalization mechanisms (59)(60) macropinocytosis contributes to TLR4 internalization in macrophages. LPS transiently increases membrane ruffling in human MDMs(61) and increases dextran uptake in mouse macrophages(23,62). Moreover, the internalization of LPS-activated TLR4 requires phospholipase C-γ2 (PLCγ2)(62), a regulator of both constitutive and inducible macropinocytosis(11,63). We therefore asked whether GAS7, by regulating macropinocytosis, modulates the two waves of TLR4 signalling. At 16h post-LPS stimulation, GAS7-silenced MDMs secreted more CXCL8 and IL6 than controls (Fig. 6c). Next, we employed qPCR to simultaneously measure the induction of pro-inflammatory mediators and ISGs at early and late time-points after LPS stimulation. At 2h post-stimulation, the induction of *CXCL8* was already elevated in GAS7-silenced MDMs in comparison with shControl cells, whereas ISGs (*RSAD2*, *MX1*, *IFI44L*) were not yet induced (Fig. 6d, S6c). At 8h post-stimulation, ISG transcripts were significantly decreased in GAS7-silenced cells (Fig. 6d, S6c), indicating defective TRIF-dependent signalling, while *CXCL8* remained elevated. PI3K inhibition with LY294002, abolished ISG induction and further increased *CXCL8* at 8h (Fig. 6d, S6c), supporting a macropinocytic internalization route for LPS-activated TLR4 in human MDMs.

Finally, re-expressing an shRNA-resistant GAS7c transgene under the *PGK* promoter in shGAS7 MDMS (Fig. S6c) restored ISG induction at 8h post-LPS stimulation, while lowering *CXCL8* transcripts (Fig. 6d, S6c). Thus, GAS7 contributes to the duality of TLR4 signalling, promoting TLR4 internalization and ISG induction, while reducing plasma-membrane NF-kB-mediated inflammation.

Collectively, our data identifies GAS7 as a F-BAR adaptor enriched in macrophages that promotes membrane ruffling-driven macropinocytosis and immune surveillance over a filopodial, motile and invasive state (Fig. S7).

## Discussion

Tissue surveillance requires macrophages to continuously sample their environment, an activity often viewed as incompatible with high motility (19). Here we identify GAS7 as a key determinant of this balance. We show that GAS7 expression in macrophages promotes constitutive fluid-phase uptake while limiting cell motility, thereby enhancing their sentinel capacity to detect and respond to microbial ligands.

Macrophages and dendritic cells perform constitutive macropinocytosis, and this flux can be further enhanced by growth factors, cytokines or microbial stimuli, such as LPS or M-CSF (3). GAS7 is required for both constitutive and induced macropinocytosis, indicating that it acts at a stage where the upstream triggers of macropinocytosis converge into a common pathway. GAS7 is enriched at membrane ruffles and required for optimal ruffle formation. It is largely absent from EEA1⁺ and LAMP1⁺ compartments, pointing to a role at early stages of macropinocytosis rather than during macropinosome maturation. As with other F-BAR proteins, GAS7 associates with membranes in a curvature-dependent manner(64). Because of its unusually shallow F-BAR domain, GAS7 binds preferentially to less curved membranes(27), where it assembles as multimeric sheet-like structures driven by lateral interactions between F-BAR dimers(27,65). In mouse macrophages performing phagocytosis, GAS7 localizes to the relatively flat base of phagocytic cups (27). This suggests that, in both macropinocytosis and phagocytosis, GAS7 assembles at low curvature regions of the plasma membrane, where it functions as a scaffold for actin regulators. As membrane curvature increases after closure of macropinocytic or phagocytic cups, GAS7 disengages from the membrane, allowing its redistribution to new sites.

The formation of macropinocytic cups depends on Arp2/3-mediated branched actin polymerization at the plasma membrane, driven by the WRC, itself activated by binding of GTP-RAC1(3,13). Our results show that GAS7 directly interacts with two subunits of the WRC, WAVE-2 and ABI-1, and is required to sustain elevated levels of active GTP-bound RAC1 in macrophages. How GAS7 sustains RAC1 activation remains unclear. Because it does not directly interact with RAC1, it is unlikely to act as a classical GEF. Instead, a plausible model is that GAS7 promotes clustering of the WRC underneath the membrane(66) This would favour local RAC1 engagement and shield RAC1-GTP from inactivation by GTPase-activating proteins (GAPs). In this way, GAS7 could sustain local RAC1 activity to support ruffle formation.

The successful assembly of macropinocytic cups requires alternating cycles of actin polymerization to initiate ruffle formation, followed by signal attenuation to allow folding and collapse of the macropinocytic cup (5). Recent work has highlighted CYRI proteins (CYRI-A/B) as negative regulators of RAC1-driven actin nucleation in macropinocytic cups(67). CYRI binds RAC1-GTP, shielding it away from the WRC, and thereby reducing actin nucleation to promote closure of macropinocytic cups(68,69). Altogether, this underscores the need for precise regulation of RAC1 activity to ensure that membrane protrusions progress into ruffles and further mature into cups and macropinosomes. Thus GAS7 and CYRI proteins may act in concert to provide spatiotemporal control over RAC1 during macropinocytosis. GAS7 potentiates RAC1 activity at the early stages of ruffle formation, while CYRIs dampen it in mature ruffles and cups to promote collapse and sealing.

In the absence of GAS7, macrophages adopt a motile phenotype characterized by filopodia formation, altered podosome dynamics, and induction of an ECM-remodelling transcriptional program. In a variety of cancer models, higher expression of GAS7 by tumour cells is associated with suppressed cell migration and decreased metastatic capacity(28–30). Moreover, in neuroblastoma cells, low expression of GAS7 leads to transcriptional upregulation of pathways involved in ECM remodelling(29). Along with our findings, this supports a role for GAS7 in restraining motile/invasive behaviour across diverse cell types.

Our findings position GAS7 as a nexus between macropinocytosis and motility, two cellular processes often viewed as antagonistic(19). Because the cellular pools of actin regulators are limited, it is argued that cells must allocate them either to protrusions that drive migration or to ruffles that generate macropinosomes(19,67). In this context, GAS7 biases the system towards ruffle formation and macropinocytosis. Notably, loss of CYRI-A results in excessive RAC1 activation promoting persistent lamellipodia that do not convert into membrane ruffles(69). This promotes invasiveness of sarcoma cells and reinforces the idea that disrupting macropinocytosis favours a migratory and invasive profile. Our data indicates that the absence of GAS7 leads to aberrant CDC42 activation. CDC42 is a major regulator of cell polarity, filopodia, podosome dynamics, and ECM-remodeling programs(24,25,50,52). In line with this, we found that CDC42(Q61L) enhanced macrophage spreading and induced ECM-remodelling genes, whereas a dominant-negative CDC42 mutant partially normalized gene expression in GAS7-deficient cells.

As with RAC1, GAS7 does not interact directly with CDC42, but binds its effectors, including WASP and FMNL2, suggesting that it may modulate their localization and availability to interact with CDC42. This may indirectly promote CDC42 inactivation by rendering it more susceptible to GAPs or guanine nucleotide dissociation inhibitors (GDIs). Interestingly, active CDC42 promotes the multimeric assembly of GAS7 at the membrane (65), suggesting the presence of a negative feedback loop, wherein an increase in the levels of active CDC42 is counteracted by larger assemblies of GAS7 at the membrane. Future work should explore mechanistically how GAS7 modulates the activities of RAC1 and CDC42.

The role of CDC42 in macropinocytosis has been a matter of debate (5). CDC42 is often placed alongside RAC1 as a mediator of membrane ruffling, as they share several GEFs and downstream effectors, making it difficult to isolate their individual contributions. Our data clarifies this aspect in macrophages by showing that enforced activation of CDC42 abolishes ruffle formation. Moreover, blocking CDC42 activity through expression of a dominant negative mutant partially restores membrane ruffling, although it cannot fully substitute for GAS7. This suggests that GAS7 promotes macropinocytosis by, first, restraining CDC42 activation and, second, by acting as a scaffold at the membrane to relocalize the actin machinery and coordinate RAC1 activation.

GAS7 promotes cell survival when MDMs rely on extracellular protein as an amino acid source, likely due to its ability to increase fluid uptake. Interestingly, GAS7 was initially identified as a growth-arrest–specific gene induced upon serum starvation in mouse fibroblasts (70,71). In contrast with fibroblasts, macrophages and DCs express GAS7 constitutively. This indicates that *GAS7* is a gene involved in cell survival under stress conditions and that has been co-opted by immune cells to assist in immune surveillance. GAS7 is required for optimal activation of NOD2 by MDP and modulates the early and late phases of the TLR4 response to LPS. Together with its reported role in phagocytosis (27,65), our data support a model in which GAS7 coordinates both detection and elimination of foreign threats. Whether GAS7 also protects macrophages from direct infection by pathogens remains an important question to be addressed in future work.

In immune cells, GAS7 is expressed almost exclusively by myeloid cells, while in non-hematopoietic cells, it is mainly detected in subsets of neurons, spermatids and fibroblasts (Human Protein Atlas). GAS7 can also be found in certain tumour cells (28–30), where its expression is upregulated by p53 (28) and repressed by *MYCN* (29). In contrast, in normal human macrophages, the control of GAS7 expression remains essentially unknown. Macrophages are highly dynamic cells, capable of changing their behaviour in response to environmental changes. Intravital imaging shows that many tissue-resident macrophages behave as relatively sessile sentinels at steady state(72), whereas upon infection they can rapidly switch to a highly motile mode(73,74). Moreover, during the resolution phase of an infection or injury, macrophages can acquire motile, ECM-remodelling phenotypes that support recruitment, clearance, and tissue repair(75). Given the ability of GAS7 to favour macropinocytic surveillance over motility and ECM remodelling, its expression is likely to be modulated by environmental cues across these states. Defining the pathways that regulate GAS7 expression, and how pathogens exploit or subvert GAS7-dependent programs, may open new avenues to therapeutically modulate macrophage dynamics and function.

## Materials and Methods

### Ethics statement

Plasmapheresis residues from adult healthy donors were provided by the Établissement Français du Sang (Paris, France) in accordance with the ethical guidelines issued by the Institut National de la Santé et de la Recherche Médicale (INSERM, France). French Public Health Law (art L 1121–1-1, art L 1121–1-2) does not require written consent by donors nor Institutional Review Board for non-interventional human studies.

### Cells and cell culture

Peripheral blood mononuclear cells (PBMCs) from plasmapheresis residues were separated using Ficoll-Paque (GE Healthcare), and monocytes were isolated by positive selection using CD14^+^ magnetic beads (Miltenyi Biotec # 130-097-052) and differentiated into MDMs for 10-12 days in RPMI (Gibco, Life Technologies) supplemented with 5% foetal calf serum (FCS; Gibco), 5% human serum AB (Sigma-Aldrich #H4522), 1% of 10,000 U/ml penicillin and 100 μg/ml of streptomycin (Gibco), and 50 ng/ml macrophage colony-stimulating factor (M-CSF; Miltenyi Biotec # 130-096-489). CD4 T cells were isolated from the CD14-negative fractions, discarded from CD14 enrichment, using a negative selection kit (Miltenyi Biotec # 130-096-533).

Human embryonic Kidney (HEK) 293 LTV cells (ATCC) were maintained by bi-weekly subpassage in DMEM (Gibco, ThermoFisher), supplemented with 10% FCS and 1% of 10,000 U/ml of penicillin and 100 µg/ml of streptomycin.

### Lentiviral production

Lentiviral particles were produced by transfection of HEK 293 LTV cells with 40 KDa polyethylenimine (PEI, Polyscience #24765-1)). Cells were seeded in culture flasks at a concentration of 93 000/cm2. The day after, transfections were performed with a mix of plasmids consisting of 16.67 ng/cm2 of CMV-VSVG envelope, 38ng/cm2 of packaging psPAX2, 1.9ng/cm2 of Vpr-any-Vpx and 56.67 ng/cm2 of lentiviral vector diluted in 9.33µL/cm2 of optiMEM. A list of all the plasmids employed in this study is available in Supplementary Table 1. The plasmid mixture was then complexed with 1.55µL/cm2 of polyethyleneimine (PEI, stock solution at 7.5 mM) diluted in 9.33µL/cm2 of optiMEM and added to cells. Culture medium was renewed 16 hours later, and viruses were harvested at around 48 hours after transfection, filtered through a 0.22 µm pore and stored at −80°C until use. Optionally, lentiviral preps were concentrated by overnight incubation at 4°C with 10% of a polyethylene glycol (PEG)-8000 solution (stock at 400g/L). Aggregated lentiviral particles were then pelleted by centrifugation (1h at 1500g), resuspended in MDM culture media and stored at -80 °C.

### Monocyte transduction

Ten million freshly purified CD14+ monocytes were plated in 100-mm culture dishes in M-CSF supplemented medium. At day 3 of culture, MDMs were transduced with 5 mL of non-concentrated lentiviral suspension (or 500 μL of a 10X concentrated suspension) in the presence of 8 μg/mL of protamine sulphate (Sigma-Aldrich #P4020).

Co-transductions of shRNA and the Cdc42 Q61L mutant were performed by first transducing shRNAs at day 3, as described above, with the mutant-encoding lentivirus added at day 5 of culture. A similar protocol was followed when co-transducing the shRNAs and the shRNA-resistant versions of GFP-GAS7C or GAS7C. In the case of CDC42 T17N, MDMs were first transduced with the mutant at day 3 post-seeding, with the shRNA-encoding lentivirus added at 5 days post-seeding. Finally, the co-transductions between shRNAs and Lifeact-mcherry were performed by adding simultaneously the two types of lentiviruses to the cells at day 3 of differentiation.

For transgene cassettes containing an antibiotic resistance, the selection was performed with 3 μg/mL of puromycin or 200 μg/mL of hygromycin, starting 2 days post lentiviral transductions. MDMs were then harvested at day 10-12 of differentiation, using Accutase solution (Gibco) for detachment. Cells were counted and replated for downstream experiments.

### Western blot

200 000 cells were lysed per condition using RIPA buffer (Sigma-Aldrich, R0278) in which cOmplete, EDTA-free protease inhibitors (Roche #11873580001) were added. Protein quantities were normalized using a BCA assay kit per manufacturer’s instructions (ThermoFisher #23235). 20µg of proteins were reduced in Laemmli buffer (ThermoFisher #J61337) and loaded on precast polyacrylamide (stain free or not) gels with a 4-20% gradient (BioRad). After transfer onto PVDF membranes (BioRad), membranes were blocked in [5% non-fat dry milk (w/v) in PBS-Tween-20 0.1% (v/v)] blocking buffer for 1h at room temperature. Primary antibodies (see supplementary Table 2) were diluted in antibody dilution buffer (PBS-5% BSA5%-0.01%Tween20) and incubated overnight at 4°C with gentle agitation. Membranes were then washed and incubated with species-matched secondary HRP-conjugated secondary antibodies for 1 h at room temperature. Membranes were revealed in a Bio-Rad Chemidoc after addition of HRP substrate (Bio-Rad #1705061). To reveal a second protein, membranes were first stripped with acidic solution (EMD #2504) and then the previous protocol was repeated. Densitometry quantifications were performed using the ImageLab software (Bio-Rad).

### Dextran and DQ-Red BSA uptake assays

MDMs were seeded in 24-well plates over 12-mm coverslips at a cell density of 150 000 cells/ well. Cells were pre-treated for 1 hour with Ly294002 (10 μM, Tocris #1130 ), CK666 (20 μM, Sigma-Aldrich) or chloroquine (50 μM, Sigma-Aldrich # SML0006) and exposed to 70 kDa FITC-dextran (200 μg/mL, ThermoFisher #D1822) for 30 minutes or to DQ-Red BSA (10 μg/mL, ThermoFisher #D12051) for 3 hours. Cells were fixed in 4% PFA, mounted with DAPI in Prolong-Gold mounting media (Thermofisher #P36934). Cells were imaged using a Leica SP8 confocal microscope equipped with a 20X objective. For dextran uptake, for each condition and each donor, a z-stack tile scan encompassing a surface of about 1000 X 1000 μm and containing 300 – 500 cells was acquired. For the red BSA-DQ assays, 5 z-stack fields were acquired for a total of 100-150 cells per donor per condition.

### Analysis of macrophage viability in the absence of free amino acids

Control or GAS7-shRNA-transduced MDMs were seeded overnight in 96-well plates at a cell density of 60 000 cells per well and cultured under standard culture conditions (RPMI + 10% FBS) or with RPMI without amino acids (US Biological #R90100110L), alone, or supplemented with 3% BSA (Euromedex #04-100-812-C) or 10% FBS. After 10 days of culture, cell viability was assessed using the CellTiter-Glo 3D Viability assay (Promega #G9681).

### Scanning electron microscopy

MDMs, transduced with either a control or GAS7 shRNA, were cultured on 12 mm round glass slides and fixed with 2.5% glutaraldehyde in culture media for 1h at room temperature. All samples were then washed in 1X PHEM buffer (60 mM PIPES, 25mM HEPES, 10 mM EGTA, 2 mM MgCl2, pH 7.3), postfixed in 1% osmium tetroxide for 1h, washed in water, and dehydrated in ethanol increasing series from 25% to 100%. Samples were desiccated using a critical point dryer (Leica), mounted on SEM stubs and sputtered with 20 nm gold/palladium. Macrophages were observed using a SEM IT700HR(JEOL) at 5kv with a secondary electron detector.

### Immunofluorescence

EGFP-GAS7c-transduced MDMs were seeded in 24-well plates over 12-mm coverslips at a cell density of 75 000 cells/ well. 48h later, cells were washed with 1X PBS, fixed using 4%PFA (or concentration glutaraldehyde for LAMP1 staining) and permeabilized in PBS-0.05% Saponin-0.2% BSA. For immuno-staining, primary antibodies targeting EGFP, WAVE2, RAB5, EEA1 or LAMP1 (see Supplementary Table 2 for a list of antibodies, their references and dilutions employed) was incubated overnight at 4°C. Slides were washed in permeabilization buffer and incubated for 2h at room temperature with the secondary antibody targeting primary antibodies and DAPI. For F-actin staining, cells were incubated for 2h with phalloidin-AF546 (1/400, ThermoFisher # A22283) and DAPI (1/400, BD # 564907). Slides were mounted in Prolong-Gold antifade mounting media. Cells were imaged using a Leica SP8 inverted confocal microscope equipped with a 63X oil objective. Five independent z-stack fields were acquired, for at least 5 GFP+ cells, per donor per marker staining.

### Cell sorting

5 million EGFP-GAS7C transduced MDMs were sorted in a BD FACSAria Fusion cell sorter to isolate GFP+ cells to have homogenous populations for immune-electron microscopy and western blot.

### Immuno-electron microscopy

EGFP-GAS7C macrophages were incubated with 5 nm BSA–gold particles (UMC, Utrecht University # BSAG5nm) at a concentration of 200 μg/mL, for 3 h in culture medium containing 1% FCS to load the endo-lysosomal system and then fixed with 4% paraformaldehyde in 0.1 M phosphate buffer (pH 7.4) for 2 h at room temperature. Sample preparation, ultrathin cryosectioning and immunolabelling were performed as described(76). Ultrathin sections were labelled with anti-EGFP antibodies followed by Protein A Gold-10nm (Utrecht University, The Netherlands). Sections were examined using a Tecnai Spirit electron microscope (FEI Company) equipped with a Quemesa digital camera (SIS).

### Membrane ruffling and podosome analysis

shControl and shGAS7 MDMs were detached, counted, and 500 000 MDMs were stained with fixable cell dyes CellTrace Violet stain (ThermoFisher #C34557) or CellTrace FarRed cell stain (ThermoFisher #C34564 A), respectively, for 30 minutes in PBS. For each donor, a second experiment was performed with the cytoplasmic dyes reversed. shControl and shGAS7 MDMs were mixed with a 1:1 ratio and 75 000 cells and plated over 12-mm coverslips. In the case of CDC42 mutants each condition was plated in a different well. Actin was stained using phalloidin as previously described and images were acquired using a Leica SP8 confocal microscope. For each donor, five z-stack fields were acquired with shcontrol stained in violet and shGAS7 stained in FarRed. Five additional fields in which cell dyes were reversed, were acquired, for a total of 30-90 cells, per donor per condition. Cells were imaged using a Leica SP8 inverted confocal microscope equipped with a 63X oil objective for a total of 30-90 cells, per donor per condition.

### Live Imaging

To live image EGFP-GAS7C-expressing MDMs (Video S1), 100 000 cells were plated in a glass bottom fluorodish (Ibidi #81218). Three to ten z-slices were acquired every 10-30 seconds, and for up to 45 minutes, on an inverted, 2-photon, Leica SP8 confocal microscope, equipped with a stage incubator maintaining cells at 37°C, 5% CO_2_ and 90% relative humidity. For the live imaging of MDMs expressing an shGAS7 and complemented with shResistant EGFP-GAS7C, we used an inverted Leica SP8 confocal microscope, equipped with a stage incubator and similar acquisition parameters as above. To monitor live macropinosome formation, 250µg/mL of dextran cf-633 (Biotium #80141) was added to the medium at the time of acquisition start.

To assess podosome dynamics, MDMs co-transduced with Lifeact-mCherry, and either control or GAS7 shRNA, were seeded overnight in glass bottom 8-well µ-Slides (Ibidi #80807) at a cell density of 50,000 cells per well. To increase the number of acquired cells, we also included MDMs additionally co-transduced with Gag-iGFP from HIV-1, from an unrelated project, which had no impact on podosome dynamics. Live imaging was performed in a Leica SP8 inverted confocal microscope, equipped with a stage incubator maintaining cells at 37°C, 5% CO_2_ and 90% relative humidity. Z-stacks encompassing the whole cell were acquired with a 63X oil objective at an acquisition rate of 1 image per 35 seconds for a total duration of 30 minutes. Videos were processed and mounted using Fiji or Imaris.

### Cell tracking

MDMs, transduced with either control or GAS7 shRNA, were stained with CMPTX CellTracker Red (Thermofisher #C34552) or CellTrace CFSE (Thermofisher #C34570), respectively (Cell trackers were reversed for the experiment shown in Fig S3e-i). One thousand cells from each condition were seeded together in 48-well plates, allowed to adhere, and left untreated or treated with 50 ng/mL of human recombinant CCL2 (Thermofisher #300-04-20UG). Cells were place in an Incucyte S3 live imaging system (Sartorius) equipped a 10X objective and green and red detection filters and imaged for 5 days at an acquisition rate of 1 image per hour.

### Image analysis

All images were analysed using Fiji version 1.54p and standardized with macro scripts which are available upon request.

Analysis of FITC-dextran and DQ-Red BSA uptake. A sum projection was generated from acquired z-stacks. The DAPI signal was thresholded to generate a mask and obtain the number of cells in each image. The total integrated intensity for the Red BSA-DQ or FITC-Dextran signal across all the field was measured and the fluorescence intensity per cell was calculated as the ratio between the total integrated intensity and the number of nuclei.

Colocalization between EGFP-GAS7C and F-actin. A binary mask of the cell (CellMask) was generated from the GFP signal at each z-slice. Based on this mask, a three-dimensional distance map of the cell was computed, in which each pixel was assigned a value corresponding to its distance to the cell boundary. A second mask defining the cytoplasmic region was then generated from this distance map, using a threshold corresponding to a cortex thickness of 1.1 µm. To isolate the cortex, the mask of the cell without the cortex was subtracted from the complete cell mask. The cortical region was then segmented into bottom and top cortexes. This segmentation was performed by measuring the cortical area for each slice, starting from the basal side, until the measured area began to decrease. The first slice at which this occurred was considered the first slice of the top cortex. The cytoplasmic mask was obtained by subtracting the cortex and nuclear masks (the latter defined from the DAPI signal) from the whole-cell mask. To verify the accuracy of mask generation, a 3D reconstruction of each cell was created, displaying each mask as an independent channel, and a 3D projection was generated using the *Reslice* tool (See Fig.2B) Finally, in a second macro, the *Coloc 2* plugin was run within each mask to calculate Pearson’s correlation coefficients between the two fluorescence channels in each area of each cell.

Colocalization analysis between EGFP-GAS7C and markers of the endocytic pathway. The GFP signal and DAPI signal were used to generate a binary mask of the cell and the nucleus respectively at each z-slice. Subtracting the nuclear mask from the cell mask provided the cytoplasm mask. The plugin coloc-2 was then run on the cytoplasm to calculate Pearson’s correlation coefficients between the two channels of interest for each cell. To generate intensity plots of stainings, a line was manually drawn across cells of interest and intensity profiles along the line for each channel were measured using the plot profile function. The intensity is represented as a percentage of the maximum intensity along the line and smoothed with the 8 (for actin staining) or 10 (for colocalization analysis) closest neighbours.

Assessment of membrane ruffles and podosome distribution. The contour of each individual cell was manually drawn using the maximum intensity Z-projection of the phalloidin channel to create a region of interest (ROI) defining the cell. Images from each condition were shuffled and images of individual cells were displayed one-by-one to 3 different volunteers blinded to cell identity. Cells were manually classified into five categories of ruffle magnitude, as revealed by the phalloidin staining: “hyperdense ruffles”, “dense ruffles”, “intermediate ruffles”, “filopodia-like” and “no ruffles”. Examples of cells classified in each category are provided in Figure S3A. Conversely, two categories of podosome distribution were defined: “homogenous” and “heterogenous” (See Figure S3B). Cell spreading was obtained from the area of each ROI.

Analysis of podosome dynamics. Maximum intensity projections of the lifeact-mCherry signal were obtained for each time frame. Subsequently, lateral drift was corrected using the *StackReg* plugin and the *Rigid Body* algorithm(77). Two masks were then generated from the lifeact signal. One with a low threshold to define the cell ventral surface and cell contour and a second, more stringent, to define podosomes. Thresholds were user-defined but applied constantly across all cells acquired. To measure podosome dynamics, we quantified the cumulative podosome coverage at time *t* (*CC_t_*), which we defined as the fraction of the ventral surface of the cell that has been visited (covered) by podosomes up to time=*t*. Thus, at each time point, areas covered by podosomes at the ventral surface that have not been previously covered are added to *CC_(t-1)_* to obtain *CC_t_*. This can be visualized on the cumulative podosome mask (Video S6) that expands as podosomes disassemble and reassemble on previously uncovered areas of the ventral surface.

We normalize the *CC_t_* at the first time-point to 1 to account for different initial podosome levels across different cells and plot its values across all the acquisition time for representative cells in Fig 3G. Donor means in Fig 3H are for *CC_t_* values at the end of the acquisition.

Cell tracking. Time-stacks containing 3 channels (green, red, phase) were exported from the Incucyte and loaded in Fiji. Bleach correction was applied to the green and red fluorescent channels by *Histogram Matching*. Lateral drift was corrected using the *Correct 3D Drift* plugin and using the phase channel as reference. Next, we thresholded the red and green channels to obtain cell masks, and employed the *Trackmate* plugin for cell tracking (78), sequentially for red and green cells. To detect spots (cells) we used the *Label Image Detector* algorithm, with an initial filter based on spot quality with default settings. The *Simple LAT Tracker* was employed to track spots overtime, using a linking max distance of 70 pixels, a gap closing max distance of 70 pixels, and gap closing max frame of 1. Tracks comprising less than 10 timeframes were excluded by filtering, and tracks consisting of cells that incorporated both red and green dies were manually excluded. Cell tracks were plotted as starting at the graph origin in μm (1 pixel = 1.24 μm) and cell speed was calculated from the Eucledian distance between two consecutive points.

### Bulk RNA sequencing

RNAseq was performed by the Next Generation Sequencing Platform at Institut Curie. RNA was isolated from shControl or shGAS7-transduced MDMs from 6 independent donors using the Nucleospin RNA kit (Macherey-Nagel #740955.50). RNA sequencing libraries were prepared from 600ng of total RNA using the Illumina® Stranded mRNA Prep Ligation library preparation kit, which allows to perform a strand specific sequencing. This protocol includes a first step of polyA selection using magnetic beads to focus sequencing on polyadenylated transcripts. After RNA fragmentation, cDNA synthesis was then performed and resulting fragments were used for dA-tailing followed by ligation of RNA Index Anchors. PCR amplification with indexed primers (IDT for Illumina RNA UD Indexes) was finally achieved with 13 cycles, to generate the final cDNA libraries. Individual library quantification and quality assessment was performed using Qubit fluorometric assay (Invitrogen) with dsDNA HS (High Sensitivity) Assay Kit and LabChip GX Touch using a High Sensitivity DNA chip (Perkin Elmer). Libraries were then equimolarly pooled and quantified by qPCR using the KAPA library quantification kit (Roche). Sequencing was carried out on the NovaSeq 6000 instrument from Illumina using paired-end 2 × 100 bp, to obtain around 30 million clusters (60 million raw paired-end reads) per sample.

### RNA-seq data analysis

RNA-Seq data analysis was performed by GenoSplice technology (www.genosplice.com). Quality control of sequencing data, analysis of reads repartition (e.g., for potential ribosomal contamination), inner distance size estimation, genebody coverage, strand-specificity of library were performed using FastQC v0.11.2, Picard-Tools v1.119, Samtools v1.0, and RSeQC v2.3.9. Reads were mapped using STAR v2.7.5a(79) on the human hg38 genome assembly and read count was performed using featureCount from SubRead v1.5.0 and the Human FAST DB v2021_3 annotations.

Gene expression was estimated as described previously (80). Only genes expressed in at least one of the two compared conditions and covered by enough uniquely mapped reads were further analyzed. Genes were considered as expressed if their FPKM value was greater than FPKM of 98% of the intergenic regions (background). At least 50% of uniquely mapped reads was required. Analysis at the gene level was performed using DESeq2(81), with samples paired by donor, using the Wald test and Benjamini-Hochberg correction. Genes were considered differentially expressed for fold-changes ≥ 1.5 and adjusted p-values ≤ 0.05. Pathway enrichment analyses and GSEA analysis(82) were performed using WebGestalt v0.4.4 (83) merging results from up-regulated and down-regulated genes only, as well as all regulated genes. Pathways were considered significant with p-values ≤ 0.05.

### Yeast-two-hybrid screen

Yeast two-hybrid screening was performed by Hybrigenics Services (S.A.S., Evry, France, http://www.hybrigenics-services.com).

The coding sequence for full-length GAS7C (NM_201433.2) was PCR-amplified and cloned into pB66 as a C-terminal fusion to the Gal4 DNA-binding domain (Gal4-GAS7). The constructs were checked by sequencing the inserts and used as baits to screen a random-primed human macrophages cDNA library constructed into pP6. pB66 derives from the original pAS2ΔΔ vector (84) and pP6 is based on the pGADGH plasmid (85).

For the GAS7C bait, 34 million clones (3-fold the complexity of the library) were screened using a mating approach with YHGX13 (Y187 ade2-101::loxP-kanMX-loxP, matα) and CG1945 (mata) yeast strains as previously described. 264 His+ colonies were selected on a medium lacking tryptophan, leucine and histidine and supplemented with 1 mM 3-aminotriazole to handle bait autoactivation. The prey fragments of the positive clones were amplified by PCR and sequenced at their 5’ and 3’ junctions. The resulting sequences were used to identify the corresponding interacting proteins in the GenBank database (NCBI) using a fully automated procedure. A confidence score (PBS, for Predicted Biological Score) was attributed to each interaction as previously described (86). The PBS relies on two different levels of analysis. Firstly, a local score considers the redundancy and independency of prey fragments, as well as the distribution of reading frames and stop codons in overlapping fragments. Secondly, a global score considers the interactions found in all the screens performed at Hybrigenics using the same library. This global score represents the probability of an interaction being nonspecific. For practical use, the scores were divided into four categories, from A (highest confidence) to D (lowest confidence). A fifth category (E) specifically flags interactions involving highly connected prey domains previously found several times in screens performed on libraries derived from the same organism. Finally, several of these highly connected domains have been confirmed as false positives of the technique and are now tagged as F. The PBS scores have been shown to positively correlate with the biological significance of interactions (87,88).

### Active RAC1 and CDC42 pull-down assays

Pull-downs of RAC1-GTP and CDC42-GTP were performed using the Active RAC1 Pull-Down and Detection Kit (ThermoFisher #16118) and the Active CDC42 Pull-Down and Detection kit (ThermoFisher #16119), respectively, and following the manufacturers’ instructions with minor adjustments. Briefly, for Rac1-GTP pull-downs, 4.0 × 10^6^ MDMs, transduced with either shControl or shGAS7, were lysed in 500 μL of the kit’s lysis/wash buffer (25mM Tris•HCl, pH 7.2, 150mM NaCl, 5mM MgCl2, 1%NP-40 and 5% glycerol). Lysates were cleared by centrifugation and 50 μL was set aside as input. The rest of the lysate was incubated with 20 μg of GST-PAK1-PDB, on a spin cup containing a glutathione resin, for 2 hours at 4°C with constant rotation. The resin was washed 3 times, and the elution was performed with 50 μL of Laemmli buffer 2X. Input and pull-down samples were analysed by western blot. For Cdc42-GTP pull-down, we found that the kit lysis buffer could not extract detectable amounts of the active GTPase, and as such we employed RIPA buffer (Sigma-Aldrich #R0278) for lysis. The following steps of the protocol were similar to the Rac1 pull-down.

### Quantitative PCR

RNA was isolated from shControl or shGAS7-transduced MDMs using the Nucleospin RNA kit (Macherey-Nagel #740955.50) and cDNA was synthesized from 1 μg of RNA using the High-Capacity cDNA synthesis Kit (ThermoFisher # 4374967). QPCR was performed in 384-well plates in a Roche Lightcycler 480 system, using TaqMan technology, with 1 μL of cDNA, 5 μL of TaqMan Universal PCR Master Mix (ThermoFisher #4304437) and 0.5 μL of a 20X mix of upstream primer, downstream primer and FAM-labelled MGB probe. *RPS18*, *GAPDH* and *HPRT1* were used as housekeeping genes for sample normalization. For each sample, the ΔCp between the gene of interest and the geometric mean of the Cp for the housekeeping genes was calculated and the data is presented as fold change from the control condition. Supplementary table 3 provides the references for the TaqMan primer/probes employed in this study.

### Quantification of cytokine and chemokine release

Quantification of secreted CXCL8, CCL2 and IL-6 shown in Figure 4f-g and Figure 6c were performed using the BD™ Cytometric Bead Array (CBA) Human IL-8, Human CCL2 and Human IL-6 Flex Sets, respectively (BD #558277, #558287 and # 558301), following the manufacturer’s instructions. The quantification of CXCL8, TNF, IL-6 and CXCL10 shown in Figure 6a-b was performed using the LEGENDplex™ Hu Anti-Virus Response Panel (BioLegend # 741270), following the manufacturer’s instructions.

### Statistical Analysis

Each data point represents an independent biological replicate derived from a distinct donor (N) unless otherwise stated. Technical replicates were averaged or the median was computed prior to statistical testing. In some experiments (eg. Fig 3d, 3i, 3k), replicates (individual cells, n) from a representative donor are shown, but their mean or median is computed before statistical analysis on multiple independent donors. Connected points indicate pairing of the same donor across experimental conditions. Parametric or non-parametric statistical tests were applied accordingly and are specified in the figure legends. Tests were two-sided, and paired analyses were used for matched donor experiments when appropriate. No data points were excluded unless objectively justified by predefined technical failure criteria. All statistical tests were computed using GraphPad Prism version 10.6.1.

### Schematic Drawings

Schematic drawings were created with BioRender.

## Data Availability

The bulk RNA-seq data generated in this study have been deposited in the NCBI Gene Expression Omnibus under accession number GSE309840. All other data supporting the findings of this study are available within the main figures and Supplementary Information. Source data are provided with this paper.

## Author Contributions

AH and VR performed and analysed most of the experiments. MSR performed electron microscopy. VH performed immunofluorescence, cell culture, biochemical analyses and analysed experiments. ST performed immunofluorescence and cell culture. FXG performed and analysed the transferrin internalization assay. AH, MM and VR developed image analysis scripts. CP performed immunofluorescence and cell culture. EG performed MDM fractionation and cell culture. AH, PB and VR designed experiments. AH, PB and VR wrote the article, which was reviewed and approved by all authors. PB and VR supervised the work, acquired funds and arranged collaborations.

## Acknowledgements

We thank Cyril Scandola from the Ultrastructural Bioimaging (Institut Pasteur) for the SEM images. We thank Mylene Bohec and Sonia Lameiras from the Next Generation Sequencing Platform (Institut Curie) for performing the bulk RNA sequencing. We thank Pierre de la Grange and Ariane Jolly from Genosplice for the bulk RNAseq analysis. We thank Susanne Reichinnek from HYBRIGENICS for assistance in the yeast-two-hybrid screen. PB and VR thank the ANRS-Maladies infectieuses émergentes (ANRS-MIE, Ref : AO 2020-1 CSS11, Ref: ECTZ191138) and SIDACTION (Ref: 20-2-AEQ-1285; ) for grants that supported this work. PB thanks ANR-10-IDEX-0001-02 PSL and LabEx DCBIOL for funding. VR thanks ANRS-MIE (Ref. 2016-1-16079) and SIDACTION (Ref 2019-FJC-12169) for individual Post-Doc Fellowships. AH thanks the Fondation pour la Recherche Médicale (FRM) and SIDACTION for individual Ph.D. fellowships. We thank Ana-Maria Lennon-Dumenil, Tala Tayoun, Ronan Thibaut, Delphine Bonhomme, Helena Izquierdo-Fernández and Andres Zucchetti (Institut Curie) for critical reading of the manuscript. We thank Tala Tayoun, Ronan Thibaut and Flavien Brouiller (Institut Curie) for assistance in cell sorting. We thank Ronan Thibaut and Delphine Bonhomme for assistance in categorizing membrane ruffles and podosomes. We thank Raphael Gauthier and Dorian Brager for isolation of PBCMs and CD14^+^ cells. We thank Tala Tayoun for assistance in drawing schematic diagrams.

