## Supplementary Figures and Legends for "GAS7 coordinates the activity of RAC1 and CDC42 to enhance macropinocytic surveillance and suppress motility in macrophages"

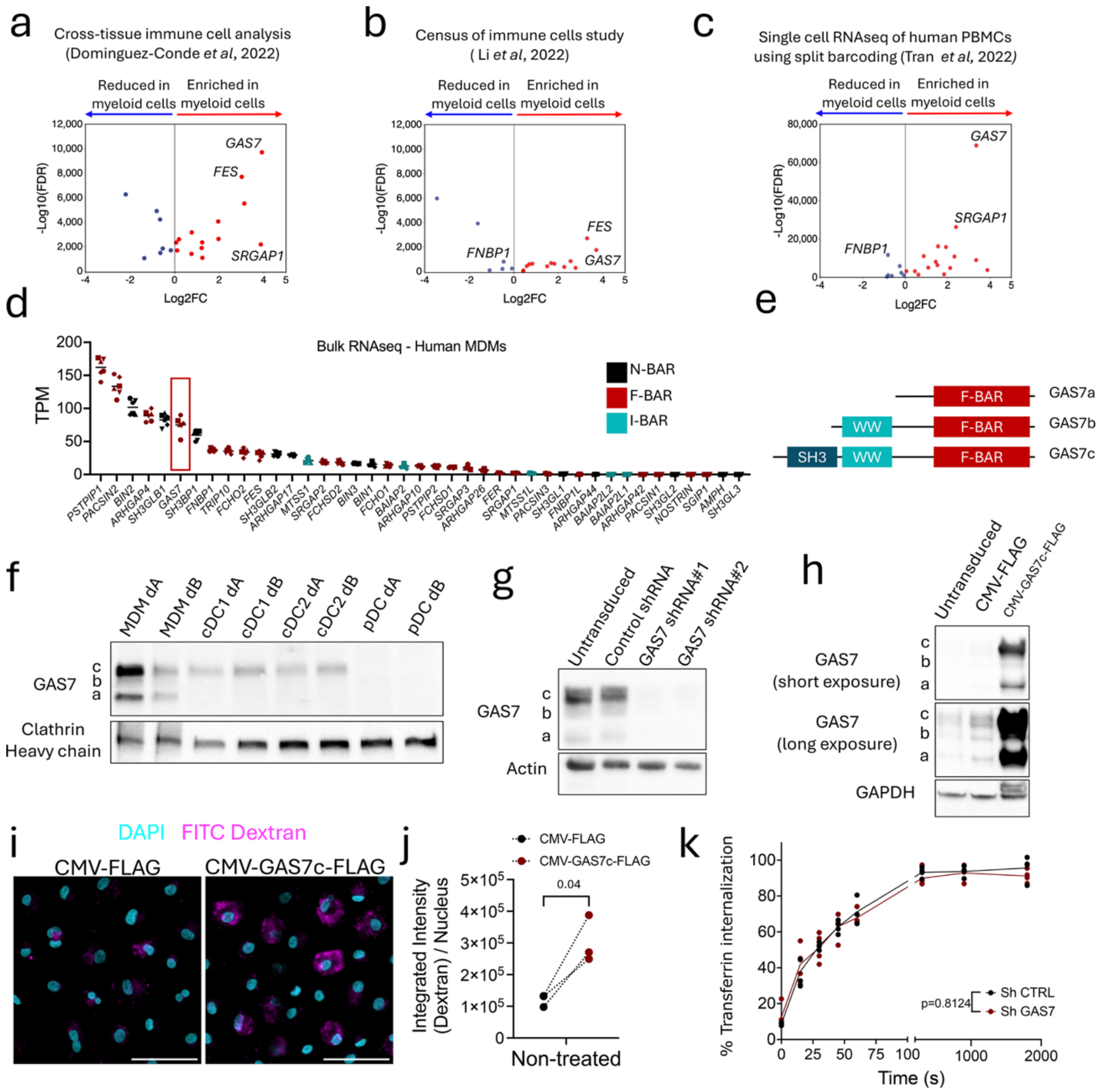


**Figure S1 – Related to Figure 1**

**a-c.** Volcano plots depicting the enrichment of BAR superfamily genes in myeloid vs non-myeloid subsets for each study indicated at the top of the plots. Genes for which cell coverage was inferior to 1% or FDR > 0.001 were filtered out and are not depicted.

**d.** BAR superfamily genes ranked by their level of gene expression in human MDMs transduced with a non-targeting shRNA, as quantified by bulk RNAseq. Independent donors are distinguished by symbol shape. TPM – Transcripts per million

**e.** Schematic depiction of human GAS7 isoforms.

**f.** Expression of GAS7 in MDMs, cDC1s, cDC2s and pDCs from two independent donors. d-donor

**g.** Western blot depicting the silencing of GAS7 in human MDMs by two independent lentiviral shRNAs

**h.** Western blot depicting the overexpression of GAS7c in MDMs via lentiviral transduction

**i.** Representative fixed-cell confocal images (SUM z-projections) of MDMs, transduced with GAS7c-FLAG or FLAG control vector, and exposed to 70 kDa FITC-Dextran (200$\mu$g/mL) for 30 min. Scale bars = 50$\mu$m.

**j.** Quantification of Fig S1I for 3 independent donors. The total integrated intensity for the FITC-Dextran channel was quantified from an acquired field and divided by the number of nuclei in the field. Each symbol represents the average dextran signal per nuclei for each donor. 300-500 cells were acquired per condition per donor. Paired t test between sh control and shGAS7 for each condition (N=3). P values depicted above brackets.

**k.** Transferrin uptake by shcontrol or shGAS7 macrophages. Data is presented as the ratio (%) between the MFI of internalized transferrin (after acid wash, to remove surface bound, but not internalized transferrin) to the total MFI. Each dot is an individual donor. Two-way ANOVA.

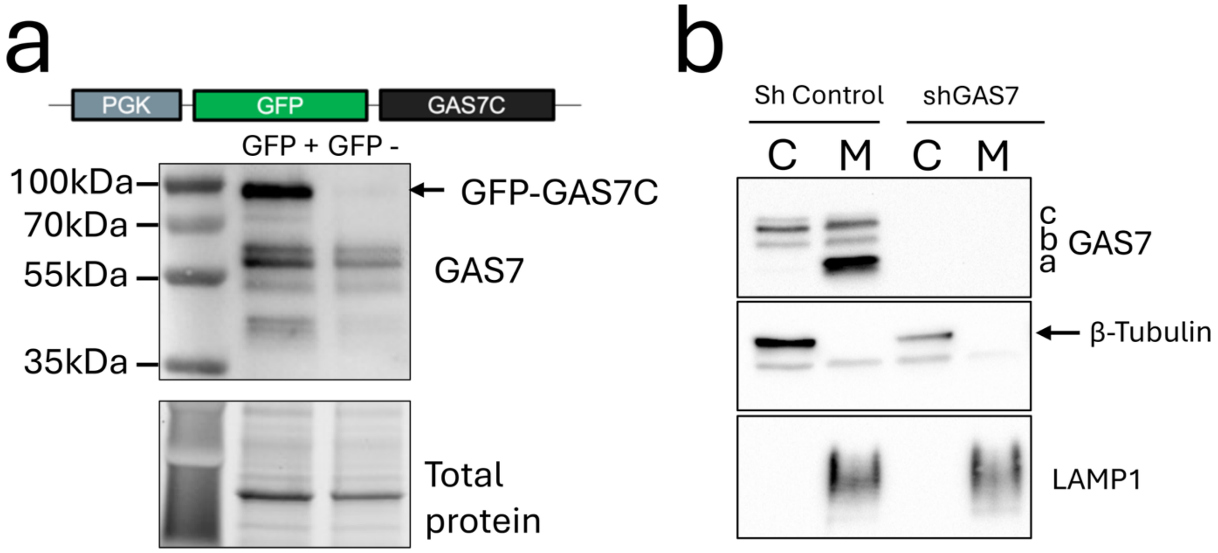


**Figure S2 – Related to Figure 2**

**a.** Schematic depiction of the construct used to express GFP-GAS7c in human MDMs via lentiviral transduction (top) and western blot of sorted GFP expressing and GFP negative MDMs showing the expression of endogenous GAS7 and the GFP-GAS7c construct (bottom).

**b.** Distribution of GAS7 in cytosolic (C) and membrane (M) fractions of macrophages in shControl or shGAS7 transduced MDMs.

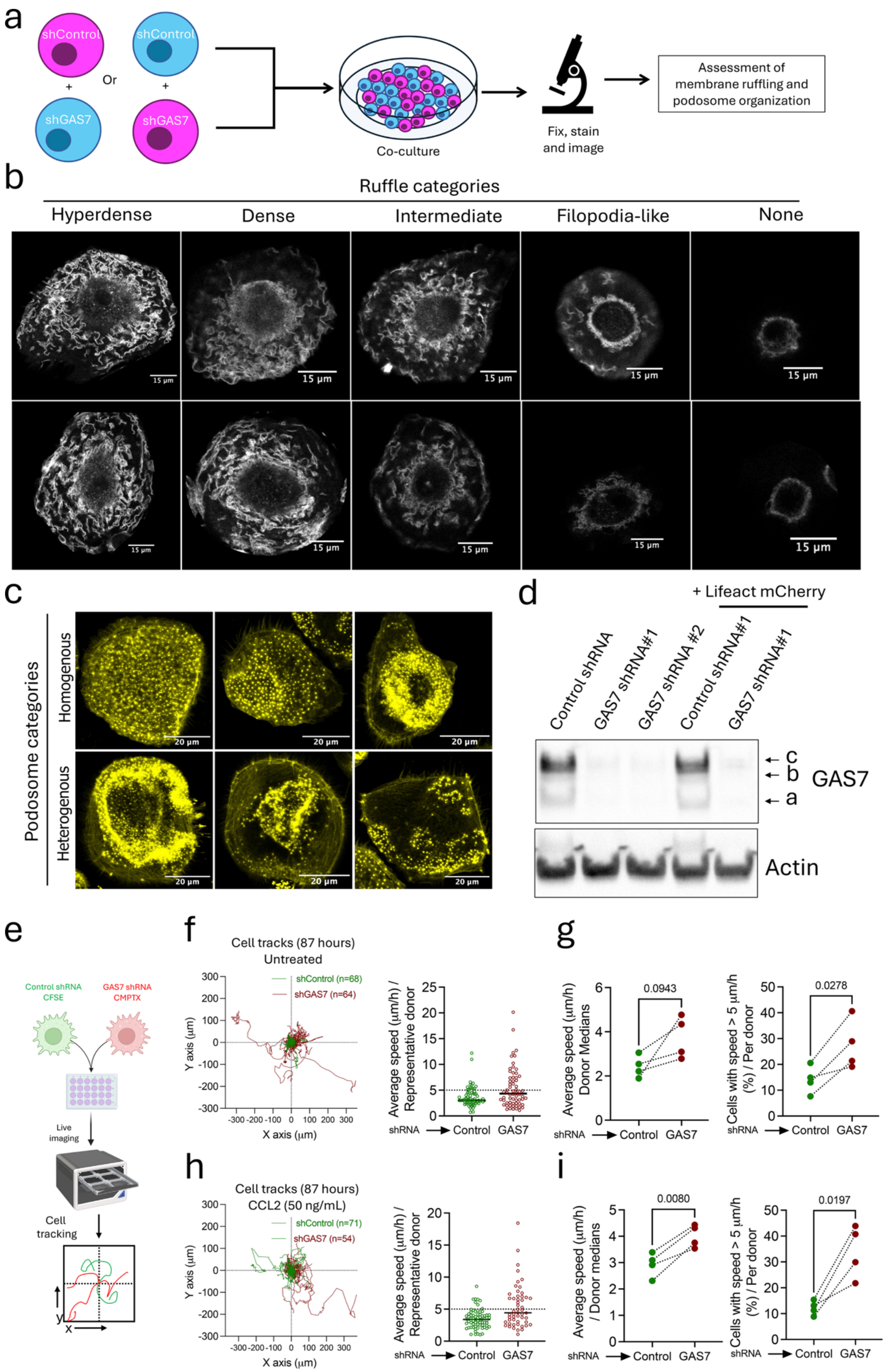


35
36 **Figure S3 – Related to Figure 3**

**a.** Schematic representation of the experimental approach used to quantify membrane ruffling. MDMs (shControl or shGAS7) were stained with Cell Trace Violet (shControl, Cyan) or Cell Trace Far Red (shGAS7, Magenta) and seeded together. Experiments with the reversed dyes were also performed and analyzed together. Cells were fixed, then subsequently stained for phalloidin, and the dorsal (top) and ventral (bottom) sections were imaged by confocal microscopy. Membrane ruffles and podosome patterning were evaluated by investigators blinded to cell identity (see Methods for details).

**b.** Representative images of cells displaying hyperdense, dense ruffles, intermediate, filopodia-like structures or no ruffles. These images were used to train investigators that performed blindly the quantifications in Fig 3B.

**c.** Representative images of podosomes considered heterogenous or homogenous. These were used to train investigators that performed blindly the quantifications in Fig 3C.

**d.** Western Blot illustrating GAS7 silencing in MDMs by lentiviral transduction, alone or in co-transduction with Lifeact-mCherry

**e.** Schematic depiction of the experimental approach used to quantify MDM movement. Cells (shControl or shGAS7) were stained with the indicated cell masks and seeded together in culture plates, in the presence or absence of CCL2 and imaged for 87 hours, at a frame acquisition rate of 1 / hour.

**f.** Trajectories (left) and average speeds (right) of individual MDMs from a representative donor in the untreated condition and pre-stained as indicated in (e), over 87 hours of live imaging. The plots were built from 2 acquisition fields, including the one displayed in Video S7. Single cells are represented by individual trajectories in (left) and symbols in (right). The black horizontal line represents the population median.

**g.** Median average speeds (left) and percentage of cells faster than 5 $\mu$m/h (right) for 4 independent donors, for the untreated condition. Each dot represents an independent donor. Paired t test.

**h.** Trajectories (left) and average speeds (right) of individual MDMs from a representative donor in CCL2-exposed cells and pre-stained as indicated in (e), over 87 hours of live imaging. The plots were built from 2 acquisition fields, including the one displayed in Video S7. Single cells are represented by individual trajectories in (left) and symbols in (right). The black horizontal line represents the population median.

**i.** Median average speeds (left) and percentage of cells faster than 5 $\mu$m/h (right) for 4 independent donors, for CCL2-treated condition. Each dot represents an independent donor. Paired t test.

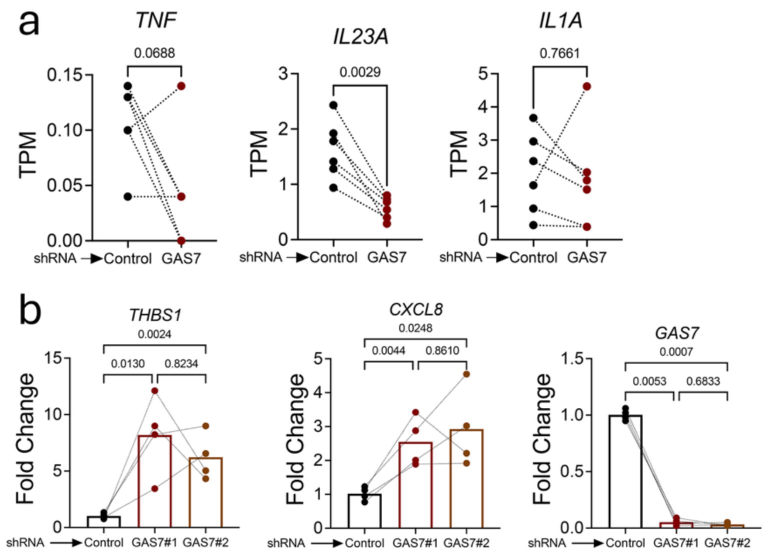


**Figure S4. Related to Figure 4**

**a.** Expression of *TNF*, *IL23A* and *IL1A* in shControl and shGAS7 MDMs in the RNAseq dataset. TPM – transcripts per million. Paired t test

**b.** Expression of *THBS1*, *CXCL8* or *GAS7* by qPCR in MDMs transduced with a control shRNA or two distincts shRNAs against GAS7. One-way ANOVA followed by Tukey's multiple comparisons test.

a

| Gene accession | Gene Name | Number of clones | PBS |
|---|---|---|---|
| NM_001178123 | ABI1 | 33 | A |
| NM_001048200 | HIPK3 | 24 | A |
| NM_005219 | DIAPH1 | 20 | A |
| NM_001258300 | KIAA1598 / SHTN1 | 18 | A |
| NM_001388182 | ATF7IP | 10 | A |
| NM_052905 | FMNL2 | 10 | A |
| NM_006990 | WASF2 | 8 | A |
| NM_201433 | GAS7 | 7 | A |
| NM_000377 | WAS | 10 | B |
| NM_213589 | RAPH1 | 4 | B |
| NM_001282799 | R3HDM1 | 3 | C |
| NM_001378957 | SF1 | 3 | C |
| NM_001386735 | ZFHX3 | 3 | C |
| XM_047432437 | FMN1 | 2 | C |
| NM_012115 | CASP8AP2 | 3 | D |
| NM_021626 | SCPEP1 | 3 | D |
| NM_001201483 | ENO1 | 1 | D |
| NM_001362972 | HMGXB4 | 1 | D |
| NM_001387869 | RAD54L2 | 1 | D |
| NM_005620 | S100A11 | 1 | D |
| NM_006842 | SF3B2 | 1 | D |
| NM_001195622 | TOPORS | 1 | D |
| NM_016312 | WBP11 | 1 | D |
| NM_001320687 | PIAS1 | 19 | E |
| NM_003345 | UBE2I | 18 | E |
| NM_001257273 | ZNF451 | 10 | E |
| NM_001410847 | HIPK1 | 6 | E |

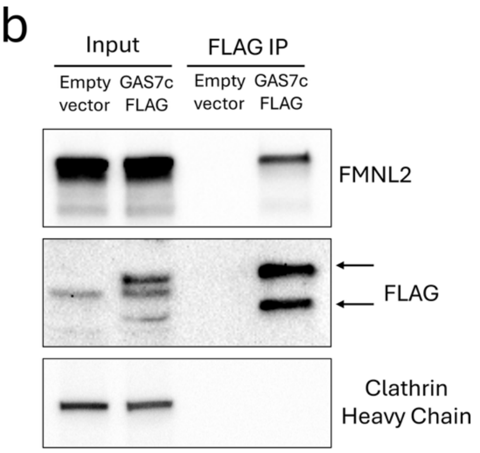


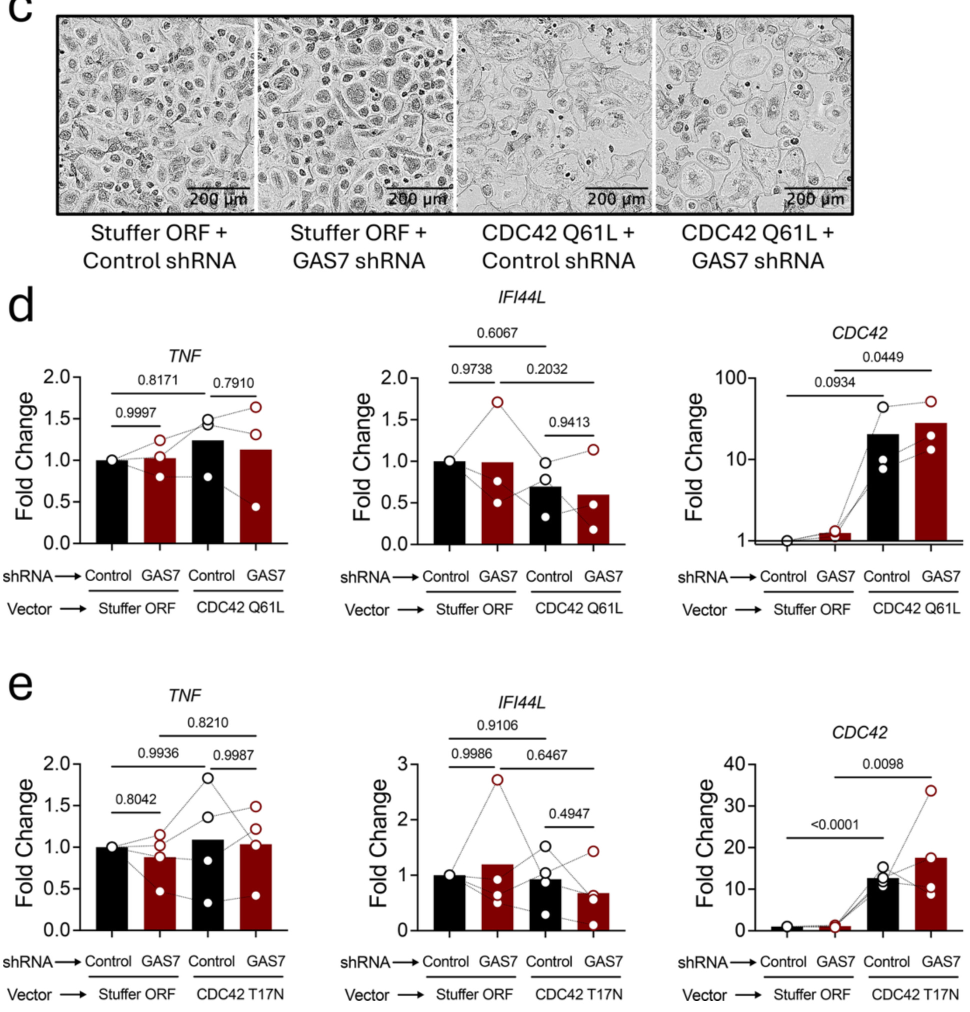


**Figure S5. Related to Figure 5**

**a.** Hits retrieved from the yeast-two-hybrid screening for binding partners of GAS7 in MDMs. The PBS (Predicted Biological Score) categorizes confidence in the interaction from very high confidence (A) to low confidence (D). Hits with scores E-F are considered as likely false positives (see Methods for details).

**b.** Co-immunoprecipitation of FMNL2 after anti-FLAG IP of in MDMs transduced with GAS7—FLAG or an empty vector.

**c.** Morphology of shControl or shGAS7 MDMs co-transduced with either a stuffer ORF or CDC42 Q61L. See also Video S8.

**d.** qPCR for *TNF*, *IFI44L* and *CDC42* in shControl or shGAS7 MDMs co-transduced with either a stuffer ORF or CDC42(Q61L). Each dot represents an independent donor and the black line connects each independent donor across the different conditions. One-way ANOVA followed by Tukey's multicomparisons test.

**e.** qPCR for *TNF*, *IFI44L* and *CDC42* in shControl or shGAS7 MDMs co-transduced with either a stuffer ORF or CDC42(T17N). Each dot represents an independent donor and the black line connects each independent donor across the different conditions. One-way ANOVA followed by Tukey's correction for multicomparisons.

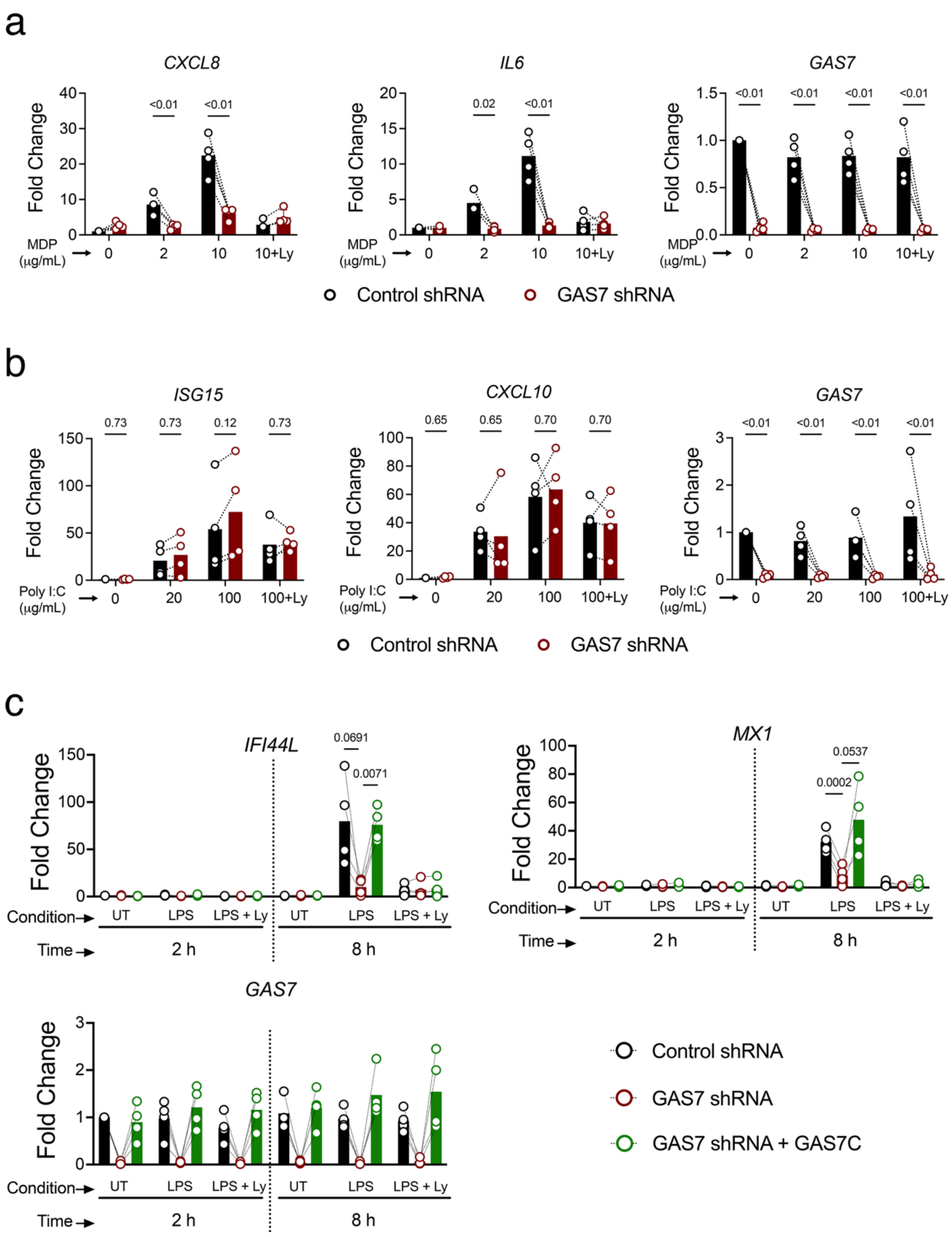


**Figure S6. Related to Figure 6**

**a.** qPCR for *CXCL8*, *IL6* and *GAS7* from shControl or shGAS7 MDMs exposed to MDP for 16 hours, at the indicated doses, and pre-treated or not with Ly294002 (10$\mu$M). Each dot represents an individual donor. Multiple paired t test between shControl and shGAS7.

**b.** qPCR of *ISG15*, *CXCL10* and *GAS7* from shControl or shGAS7 MDMs exposed to high molecular weight Poly I:C for 16 hours, at the indicated doses, and pre-treated or not with

Ly294002 (10$\mu$M). Each dot represents an individual donor. Multiple paired t test between shControl and shGAS7.

**c.** qPCR of *IFI44L*, *MX1* and *GAS7* from shControl or shGAS7 MDMs exposed to LPS for 2 or 8 hours, at 100 ng/mL, and pre-treated or not with Ly294002 (10$\mu$M). Each dot represents an independent donor and the black line connects each independent donor across the different conditions. Two-way ANOVA followed by Tukey's test for multiple comparisons.

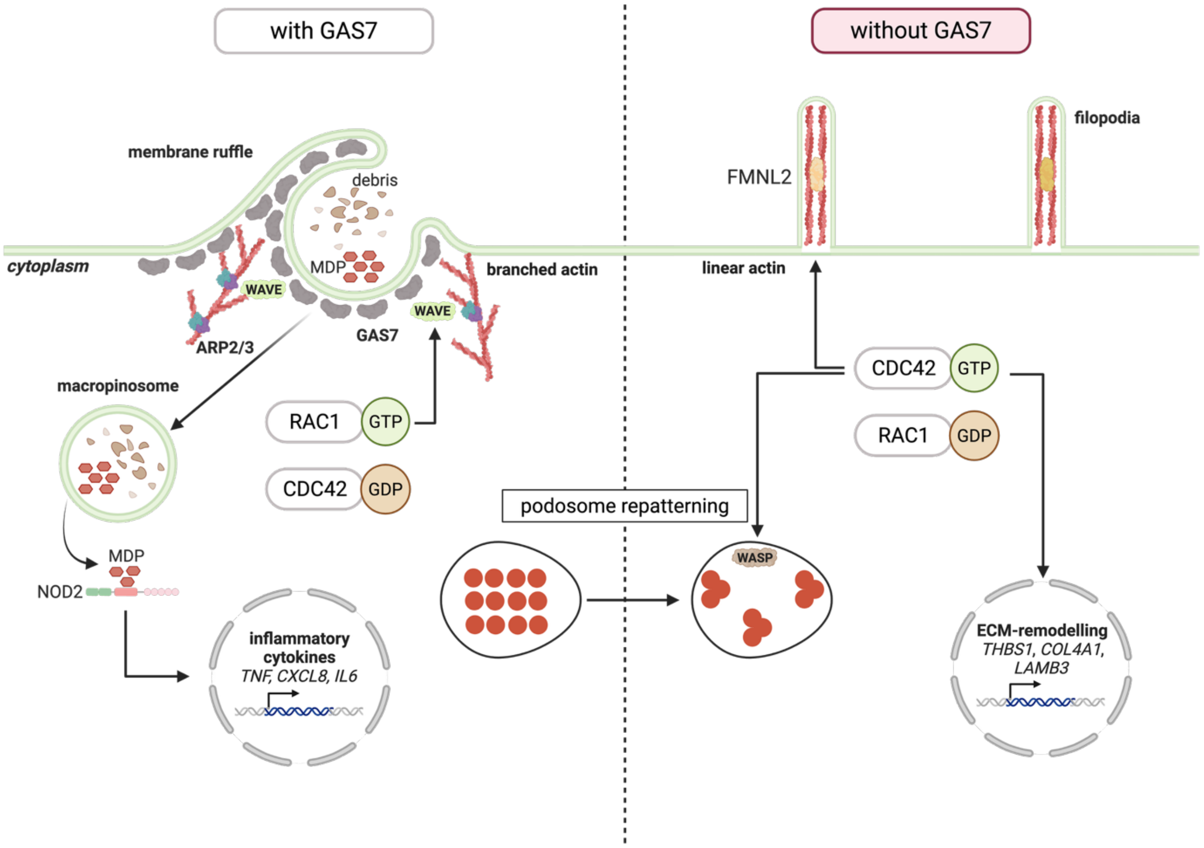


**Figure S7.** Schematic model summarizing the role of GAS7 in macrophages.

GAS7 associates with the plasma membrane, recruiting the WAVE complex and sustaining RAC1 activity to promote branched actin polymerization by the Arp2/3 complex. This enhances the capacity of macrophages to interrnalize extracellular fluid and their ability to respond to microbial ligands such as MDP. In the absence of GAS7, RAC1 activation decreases, while CDC42 becomes chronically active. This diminishes membrane ruffling, fluid uptake and innate sensing, while promoting cell movement through podosome repaterning, assembly of filopodia and production of ECM-remodelling factors.
