## Supplementary Material and Methods for "GAS7 coordinates the activity of RAC1 and CDC42 to enhance macropinocytic surveillance and suppress motility in macrophages"

### Supplementary materials and Methods

#### Identification of BAR-domain containing genes enriched in myeloid cells in humans

A list of BAR, F-BAR and I-BAR domain containing genes in humans was retrieved from the HUGO Gene Nomenclature Committee (HGCN; available at https://www.genenames.org/) for the gene groups “N-BAR”, “F-BAR” and “I-BAR”.

Bioinformatic analyses were performed using the BioTuring Platform (https://academic.bioturing.com/). To assess the relative expression of BAR superfamily genes across immune cell types we selected 3 studies that analysed the transcriptome of the major immune cell types at the single cell level, both in the blood and in tissues(31–33). To identify BAR genes enriched in myeloid cells, we employed the cell type-clusters, as annotated by the original authors of each study, to group “Myeloid” and “Non-myeloid” cells. Next, we employed the differential gene expression (DEG) analysis tool from BioTuring, and the Wilkinson test to compare the median expression levels of each BAR gene between Myeloid and Non-Myeloid groups. Genes expressed in less than 1% of the cells were filtered out. Data was displayed as a volcano plot.

#### Transferrin internalization assay

Transferrin internalization assays were performed as previously described (89). MDMs, transduced with control or GAS7 shRNAs were harvested and incubated with Tf-Alexa647 (ThermoFisher, 0.5 mg/mL) for 30 min on ice, washed, and shifted to 37°C for various periods of time in culture medium supplemented with 20 mM HEPES. After washing in PBS, half the samples were further washed in 25 mM glycine-HCl–125 mM NaCl (pH 2.8) and rapidly neutralized with 25 mM Tris (pH 10). Samples were washed one final time and analyzed by flow cytometry. The ratio of the intracellular MFI (after acid wash) to the total MFI at each time point was plotted as a function of time.

#### Subcellular fractionation

Cytosolic and membrane fractions were obtained from cultured MDMs (2 x $10^6$ cells) using the Subcellular Protein Fractionation Kit (Thermofisher #78840), following the manufacturer’s instructions. Fractions were subsequently analysed by western blot. Beta-tubulin was employed as a cytosol marker, while LAMP1 was used for the membrane fraction.

#### Cell sorting

Dendritic cell subsets were sorting as previously reported (90). Briefly, total blood DCs were enriched from human PBMCs using EasySep human pan-DC pre-enrichment kit (Stemcell Technologies #19251). The DC enriched fraction was stained with antibodies for HLA-DR-APCeFluor780, CD1c-PerCPeFluor710 (eBioscience), CD123-Viogreen, CD45RA-Vioblue (Miltenyi), Axl-PE (Clone #108724, R&D Systems), CD33-PE-CF594, Clec-9A-PE (BD) and with a cocktail of antibodies against lineage markers CD19 (Miltenyi), CD3, CD14, CD16 and CD34 (BD) in the FITC channel. pDCs were sorted as Lin$^{-}$ HLADR$^{+}$ CD33$^{-}$ CD45RA$^{+}$ CD123$^{+}$ Axl$^{-}$. cDC2 were sorted as Lin$^{-}$ HLADR$^{+}$ CD33$^{+}$ CD45RA$^{-}$ CD123$^{-}$ CD1c$^{+}$. cDC1 were sorted as Lin$^{-}$ HLADR$^{+}$ CD33$^{+}$ CD45RA$^{-}$ CD123$^{-}$ Clec9A$^{+}$. Half a million cells from each DC subset were then lysed in RIPA buffer and the expression of GAS7 was analysed by western blot.

**Immunoprecipitation**

MDMs transduced with GAS7C-FLAG or an empty lentivector (3 x $10^6$ cells) were lysed in 550 μL of NP-40 cell lysis buffer (50 mM Tris, pH 7.4, 250 mM NaCl, 5 mM EDTA,1% NP-40) supplemented with protease inhibitors. The lysate was cleared by centrifugation and incubated overnight at 4°C with constant rotation with 20 μL of pre-equilibrated magnetic beads coupled to an anti-FLAG M2 antibody (Sigma-Aldrich #M8823). Beads were washed 3 times with TBS on a magnetic stand, and elution was performed by incubating beads with 50 μL of a solution of FLAG peptide (Sigma Aldrich #F3290) at a concentration of 200 μg/mL for 30 min at room temperature with agitation. Input and IP samples were then analysed by western blot for the expression of GAS7.
